# Power and coherence in the EEG of the rat: impact of behavioral states, cortical area, lateralization and light/dark phases

**DOI:** 10.1101/2020.08.25.265520

**Authors:** Alejandra Mondino, Matías Cavelli, Joaquín Gonzalez, Lucía Osorio, Santiago Castro-Zaballa, Alicia Costa, Giancarlo Vanini, Pablo Torterolo

## Abstract

The sleep-wake cycle is constituted by three behavioral states: wakefulness (W), non-REM (NREM) and REM sleep. These states are associated with drastic changes in cognitive capacities, mostly determined by the function of the thalamo-cortical system. Thalamo-cortical activity can be examined by means of the intra-cranial electroencephalogram (iEEG).

With the purpose to study in depth the basal activity of the iEEG in adult rats, we analyzed the spectral power and coherence of the iEEG during W and sleep in the paleocortex (olfactory bulb), as well as in motor, somatosensory and visual neocortical areas. We also analyzed the laterality (right Vs. left hemispheres) of the signals, as well as the iEEG in function of the light and dark phases.

We found that the iEEG power and coherence of the whole spectrum were largely affected by behavioral states and were highly dependent on the cortical areas recorded. We also determined that there are night/day differences in power and coherence during sleep, but not in W. Finally, while we did not find right/left differences in power either in W or sleep, we observed that during REM sleep intra-hemispheric coherence differs between both hemispheres.

We conclude that the iEEG dynamics is highly dependent on the cortical area and behavioral states. We also determine that there are light/dark phases disparities in the iEEG that emerge during sleep, and that intra-hemispheric connectivity differs between both hemispheres only during REM sleep.

## 1. Introduction

The brain is a complex system, in which parallel processing coexist with serial operations within highly interconnected networks, but without a single coordinating center. This organ integrates neural events that occur at different times and locations into a unified perceptual experience (Singer, 2007; Singer, 2015). Cognitive states are mostly determined by the function of the thalamo-cortical system (Torterolo et al., 2019a). Part of this neuronal processing can be accurately measured by intra-cranial electroencephalogram (iEEG) or electro-corticogram, which avoids scalp-filtering effects that occurs mainly on the high frequencies signals with the standard surface electroencephalographic recordings.

The sleep-wake cycle is a critical physiological process and one of the most preserved biological rhythms through evolution (Torterolo et al., 2019b). This cycle is composed of wakefulness (W), non-rapid eye movement (NREM) and rapid eye movement (REM) sleep states, that are distinguished by their behavior and electrophysiological signatures, which can be captured by iEEG signals (Steriade et al., 1993; Torterolo et al., 2019a). Accompanying these electro-cortical differences among states, the cognitive capacities drastically change during the cycle. Fundamentally, consciousness is lost during deep NREM sleep, and emerges in an altered fashion during REM sleep, when most vivid dreams occur (Tononi and Laureys, 2009; Torterolo et al., 2019a).

One of the most studied neural correlates of consciousness are the cortical iEEG oscillations (Crick and Koch, 1990), which contain broad and complex frequency spectrums that can be examined by means of the fast Fourier transform. The power of the different frequencies components of the iEEG reflects the local degree of synchronization of the extracellular potential, which is deeply modified on passing from W to sleep (Buzsaki et al., 2012; Cavelli et al., 2017). While W and REM sleep contain high frequency activity together with theta waves (5-9 Hz) in the iEEG, NREM sleep shows prominent slow oscillations (delta band, 0.5 to 4 Hz) and spindles (sigma band, 11 to 15 Hz) (Achermann, 2009; Torterolo et al., 2019a; Vanderwolf, 1969; Winson, 1974).

Synchronization between oscillations from different areas represent another neural correlate of consciousness (John, 2002). In this regards, the degree of iEEG coherence between two cortical regions reflects the strength of the functional interconnections (re-entries) that occur between them (Edelman and Tononi, 2000). In other words, the spectral coherence analysis of the iEEG is a valid approach to infer cortical connectivity and communication between distant brain areas (Cantero et al., 2000). (Siegel et al., 2012) proposed that frequency-specific correlated oscillations in distributed cortical networks provide indices or ‘fingerprints’, of the network interactions that underlie cognitive processes. During W, there is a larger coherence in gamma (35-100 Hz) and high frequency oscillations (HFO, up to 200 Hz) than during sleep (Castro et al., 2013; Cavelli et al., 2015; Cavelli et al., 2017; Cavelli et al., 2018). A high degree of delta and sigma synchronization occurs during NREM sleep (Achermann and Borbely, 1998), while theta coherence is large during REM sleep in the rodent iEEG (Cavelli et al., 2018).

Although there are several studies that have analyzed the iEEG during W and sleep, we consider that a detailed and systematic evaluation of the power and coherence during these behavioral states is still pending. Hence, the purpose of this study was to provide an exhaustive examination of power and coherence of the basal iEEG activity of the adult rat during W and sleep. To this end, we studied the influence of the cortical site, employing electrodes located on the olfactory bulb (OB), frontal (primary motor or M1), parietal (primary somatosensory, S1) and occipital (secondary visual, V2) cortex, as well as the impact of laterality (differences in signals recorded in the right and left hemispheres) and the influence of dark and light phases. Upon considering this set of factors, we found important patterns of activity characterizing each sleep state along with state independent modulations of iEEG activity.

## 2. Results

Polysomnographic recordings, hypnogram and spectrogram (power spectrum as a function of time) during the light phase of a representative rat are displayed in Figure 1. As it is exhibited in this Figure, the quality of the recordings allowed an optimal classification of W and sleep epochs.

**Figure 1.**
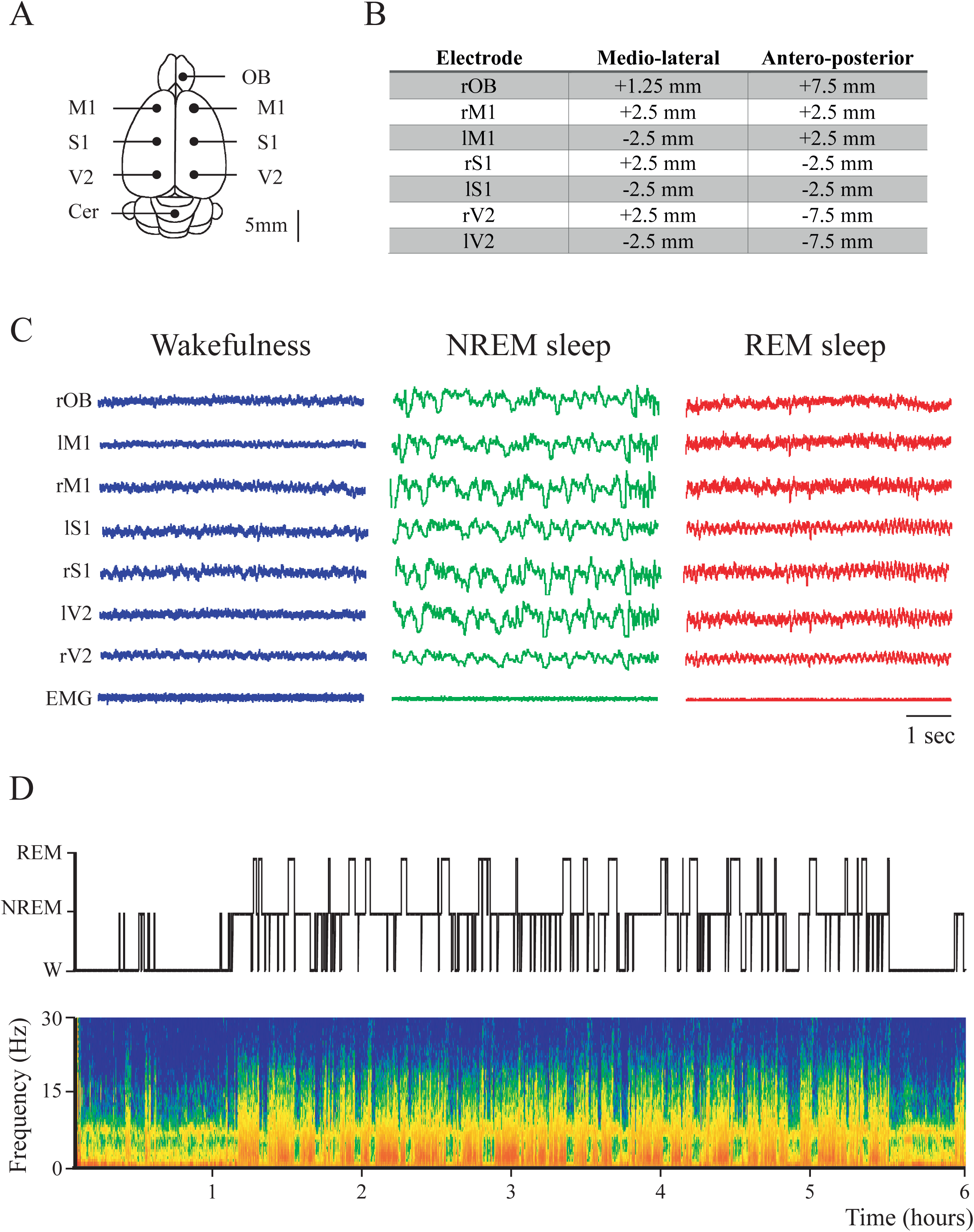
Sleep-wake states in the rat. A. Schematic representation of the electrode position in the brain of the rat. B. Electrodes’ positions in reference to *Bregma* (Paxinos and Watson, 2005). OB, olfactory bulb; M1, primary motor cortex; S1, primary somato-sensory cortex; V2, secondary visual cortex; r, right; l, left. C. Representative iEEG and the neck electromyogram (EMG) recordings during wakefulness (W, blue), NREM (green), and REM sleep (red). From top to bottom, olfactory bulb (OBr), right and left primary motor (M1r/M1l), primary somatosensory (S1r/S1l), and secondary visual (V2r/V2l) cortices. D. Hypnogram (top) according to visually scored behavioral states and spectrogram (0.1 to 30 Hz). During W and REM sleep, theta activity (5-9 Hz) in the spectrogram can be readily observed. During NREM sleep, delta activity (1-4 Hz) is more prominent and there are intermittent episodes of sigma activity (10-15 Hz) which corresponds to the presence of sleep spindles. Color calibration of the spectrogram is not provided (larger power is shown in red).

### 2.1. Power spectrum: effect of behavioral states and recording site

Figure 2 shows the absolute power spectra analysis of the iEEG during W and sleep for the OB and neocortical areas during the light (resting) phase for the right hemisphere. In order to improve the visualization of the power differences among states, in the Figure we decolorized the tracings; i.e., the pink noise or 1/f component was eliminated by multiplying the power at each frequency by the frequency itself. It is readily observed that the spectrum is highly variable in function of behavioral states and electrode sites.

**Figure 2.**
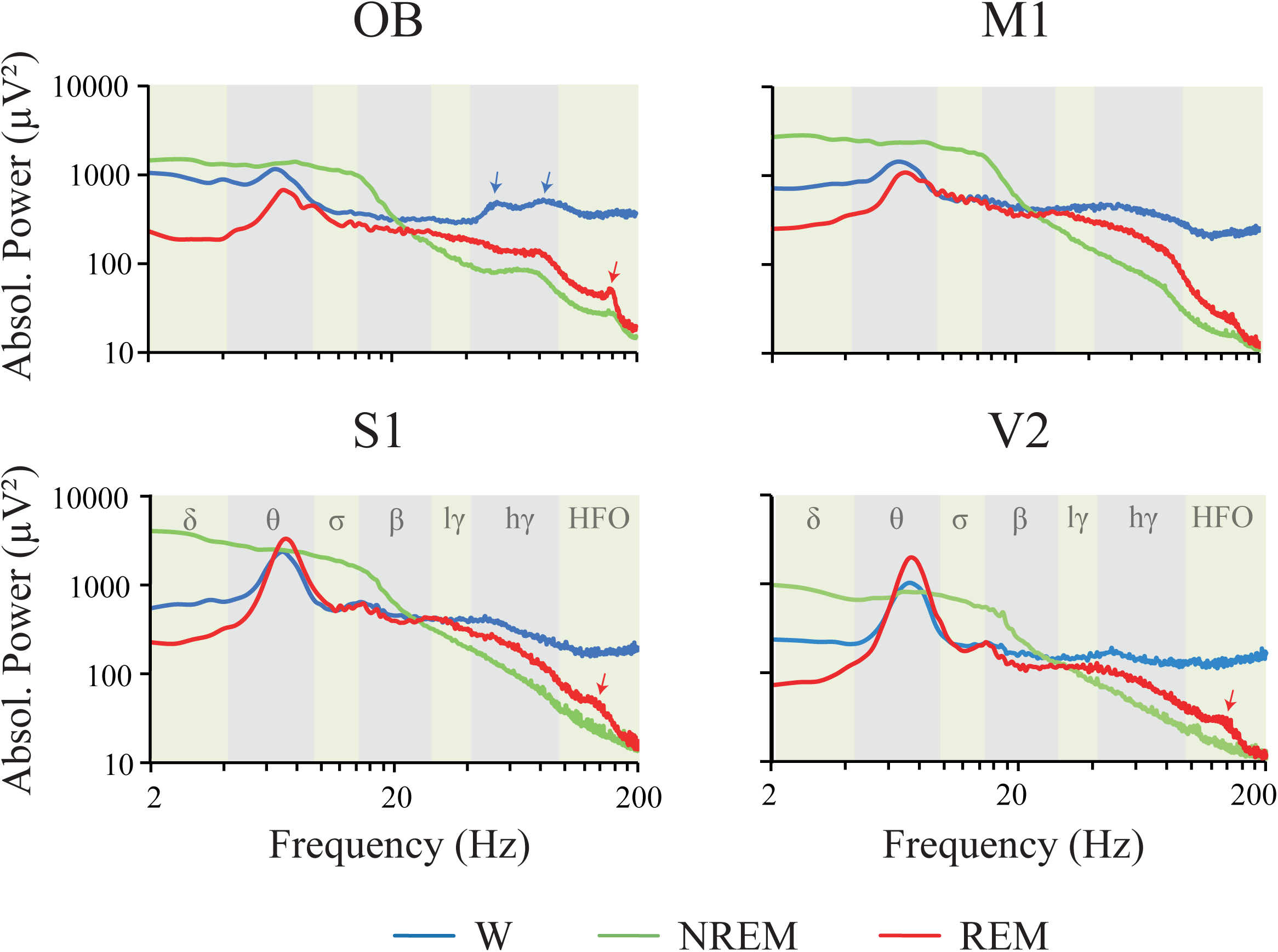
Power spectral profiles. Mean absolute power spectral profiles of the right hemisphere in wakefulness (W), NREM and REM sleep during the light period. The analyzed frequency bands are indicated by different colors in the background of the graphics. OB, olfactory bulb; M1, primary motor cortex; S1, primary somato-sensory cortex; V2, secondary visual cortex; W, wakefulness; r, right; l, left; lγ, low gamma or LG; hγ, high gamma or HG; HFO, high frequency oscillations.

The absolute power of all the frequency bands of the iEEG was affected by behavioral states, localization of electrodes and the interaction of both factors (Table 1). The exceptions were sigma, beta and low gamma (LG) that were not significantly affected by the interactions between behavioral states and electrode locations.

**Table 1.**
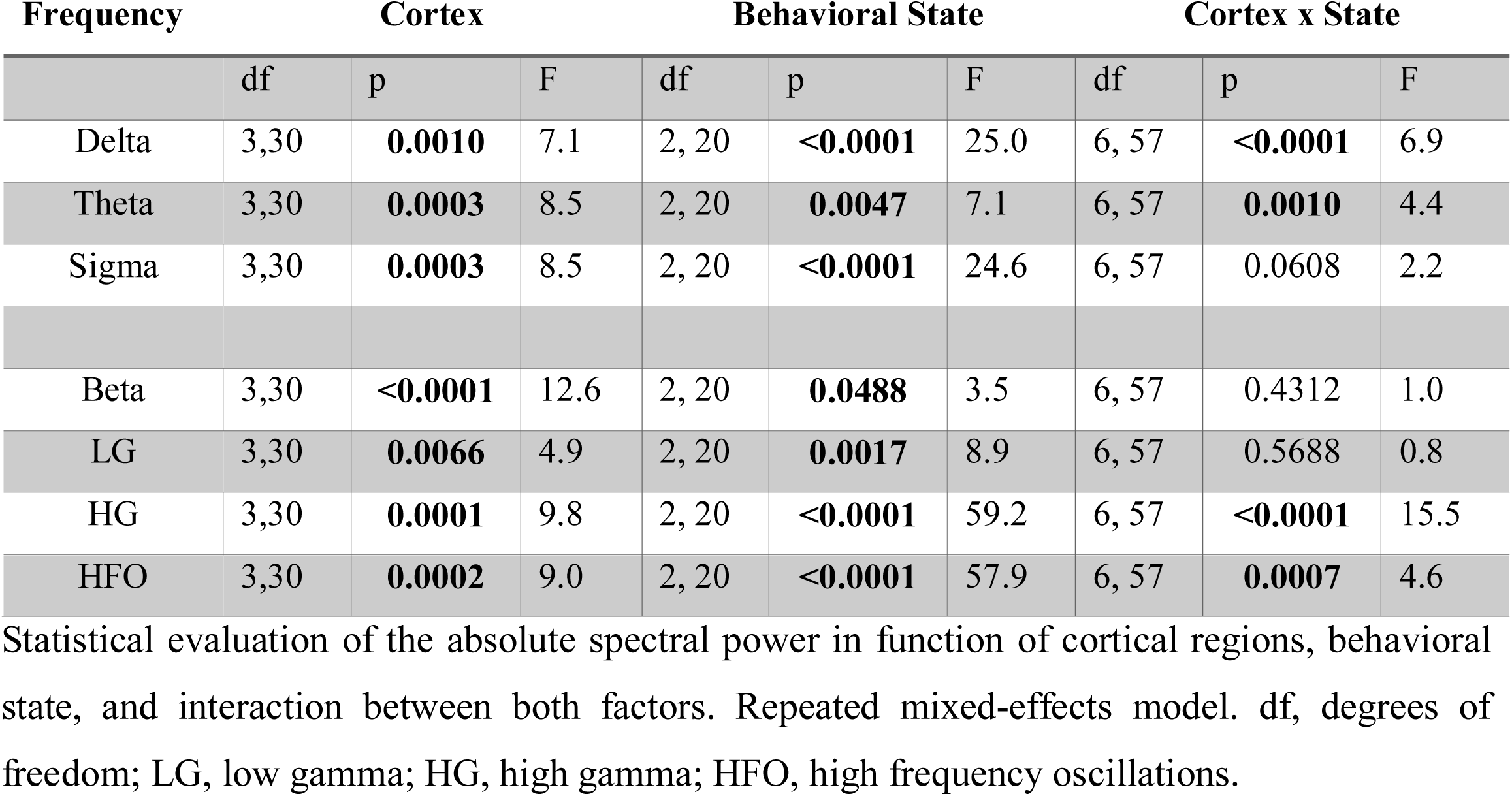
Absolute power.

A summary of the power spectrum differences between W, NREM and REM sleep is shown in Figure 3 (statistics are shown in Supplementary Figure 1A). The most remarkable results are the following. Delta, theta and sigma power during NREM sleep were significantly higher than during W and REM sleep in M1 and S1. In the OB, delta was larger during NREM compared to REM sleep, while sigma was larger during NREM compared to the other states. LG, high gamma (HG) and HFO powers were higher during W than during NREM in all the cortical areas. Also, HG and HFO during W was higher in comparison to REM sleep in most cortical areas. Finally, HG power was also larger in REM sleep than in NREM sleep in M1.

**Figure 3.**
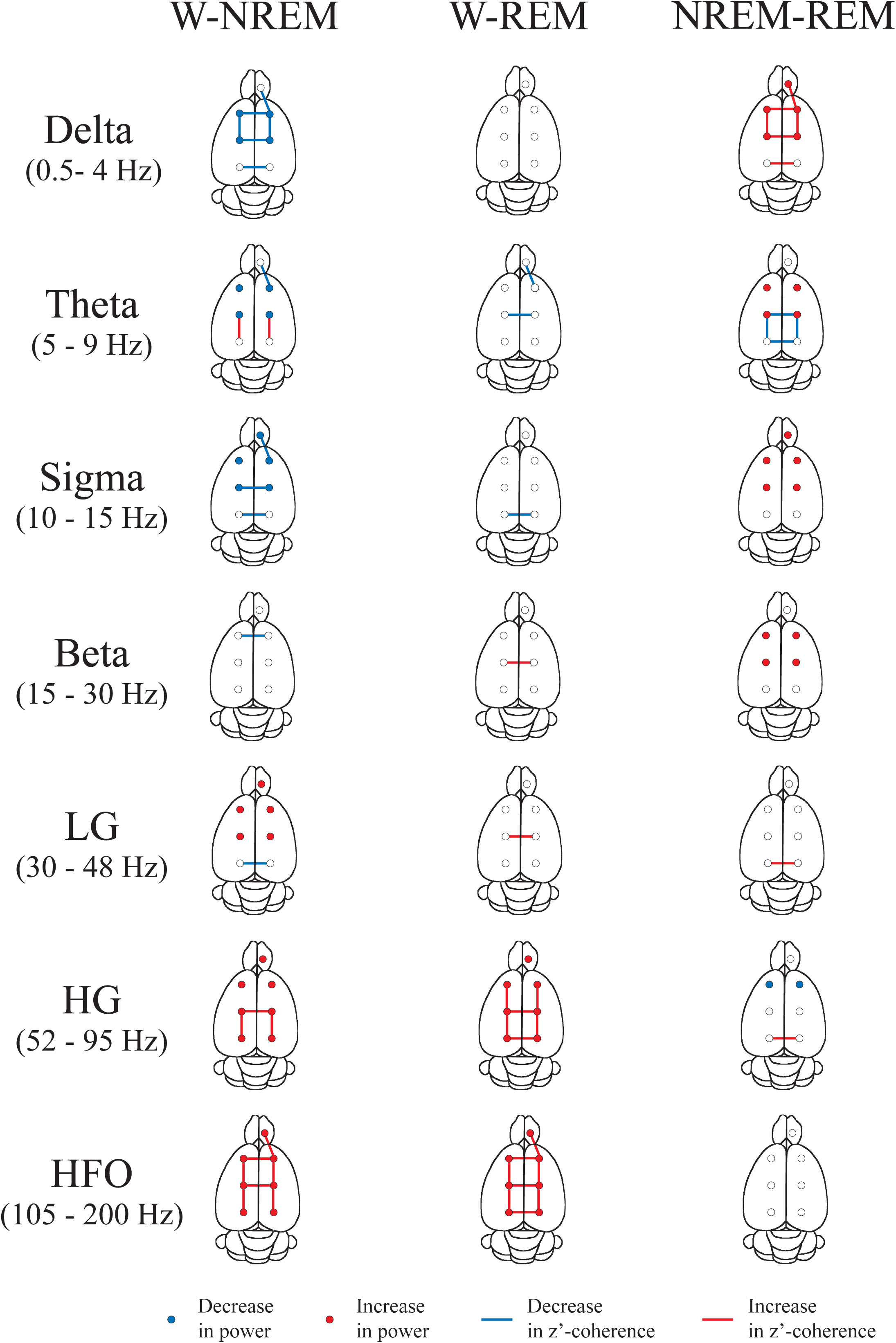
Summary of the power and z’-coherence. Statistical significant differences in absolute power and z’-coherence during wakefulness (W), NREM and REM sleep during the light phase. The circles represent the power for the different electrodes’ positions, while the lines represent the coherence for the different derivations. The results were evaluated by means of repeated measures mixed-effects model and Sidak test for multiple comparisons. Blue represents a significantly (p<0.05) lower difference between two behavioral states, and red a significantly higher difference. Power data are from the right hemisphere but are represented bilateral. The complete statistics of these data are shown in Supplementary Figure 1 and 8.

In Supplementary Figure 2, the same tracings of Figure 2 were re-plotted for a more precise comparison as function of the electrode location; the statistics of these data are shown in Supplementary Figure 1B. However, it is important to consider that the variation in the absolute power in function of the electrode site is dependent of the distance between the active and the referential electrode (cerebellum). In this regard, as shown in Supplementary Figure 3, the total power was highly modified as a function of the electrode site (total power was also affected by behavioral states and the interaction with electrode localization). Indeed, total power was the lowest in V2 (closer to the reference electrode) reaching significance during W and NREM sleep. Then, we judged to be more adequate to explore the relative instead of the absolute power in function of the electrode site (nevertheless, as we mentioned, complete statistics of the absolute power in function of the electrode site is provided in Supplementary Figure 1B). The analysis of the relative power in function of the electrode site is exhibited in Figure 4 and Table 2A and B (the analysis of the relative power in function of behavioral states is shown in Supplementary Figure 4).

**Table 2.** Relative power.

**A**
| Frequency | Cortex |  |  | Behavioral State |  |  | Cortex x State |  |  |
| --- | --- | --- | --- | --- | --- | --- | --- | --- | --- |
|  | df | p | F | df | p | F | df | p | F |
| Delta | 3,30 | 0.5710 | 0.7 | 2, 20 | **<0.0001** | 41.4 | 6, 57 | **<0.0001** | 14.9 |
| Theta | 3,30 | **<0.0001** | 35.7 | 2, 20 | **0.0009** | 10.2 | 6, 57 | **<0.0001** | 61.2 |
| Sigma | 3,30 | **0.0005** | 8.0 | 2, 20 | **<0.0001** | 15.9 | 6, 57 | 0.2353 | 1.4 |
| Beta | 3,30 | **0.0389** | 3.2 | 2, 20 | **0.0084** | 6.1 | 6, 57 | **0.0166** | 2.9 |
| LG | 3,30 | 0.0544 | 2.8 | 2, 20 | **<0.0001** | 26.8 | 6, 57 | **<0.0001** | 4.4 |
| HG | 3,30 | **0.0162** | 4.0 | 2, 20 | **<0.0001** | 22.5 | 6, 57 | **0.0213** | 2.7 |
| HFO | 3,30 | 0.0052 | 5.2 | 2, 20 | **<0.0001** | 33.6 | 6, 57 | 0.1817 | 1.5 |

**B**
| State | Comparison | Delta | Theta | Sigma | Beta | LG | HG | HFO |
| --- | --- | --- | --- | --- | --- | --- | --- | --- |
| **W** | OB vs M1 | 0.9621 | 0.8365 | 0.2279 | 0.7908 | **0.0485** | 0.9294 | **0.0334** |
|  | OB vs S1 | 0.6436 | **0.0452** | 0.0613 | **0.0274** | **0.0494** | **0.0366** | **0.0014** |
|  | OB vs V2 | 0.2194 | 0.1459 | 0.6616 | 0.7521 | 0.7853 | **0.0173** | 0.9786 |
|  | M1 vs S1 | 0.9897 | 0.4926 | 0.9954 | 0.4594 | >0.9999 | 0.3488 | 0.9120 |
|  | M1 vs V2 | 0.7637 | 0.8360 | 0.9809 | >0.9999 | 0.5783 | 0.2254 | 0.1934 |
|  | S1 vs V2 | 0.9919 | 0.9958 | 0.7595 | 0.4585 | 0.5776 | >0.9999 | **0.0137** |
| **NREM** | OB vs M1 | 0.6010 | 0.9957 | 0.2926 | 0.1330 | >0.9999 | 0.9631 | 0.9925 |
|  | OB vs S1 | **0.0054** | **<0.0001** | >0.9999 | 0.9986 | >0.9999 | 0.9711 | 0.9987 |
|  | OB vs V2 | 0.8876 | 0.09998 | 0.9974 | >0.9999 | 0.9998 | 0.9986 | >0.9999 |
|  | M1 vs S1 | 0.2591 | **<0.0001** | 0.4462 | 0.3735 | >0.9999 | >0.9999 | >0.9999 |
|  | M1 vs V2 | 0.9983 | 0.9957 | 0.6233 | 0.0928 | 0.9998 | 0.9994 | 0.9944 |
|  | S1 vs V2 | 0.0970 | **<0.0001** | 0.9999 | 0.9931 | >0.9999 | 0.9996 | 0.9991 |
| **REM** | OB vs M1 | 0.9979 | 0.7428 | **0.0018** | 0.0698 | 0.0872 | >0.9999 | 0.2567 |
|  | OB vs S1 | 0.5279 | **<0.0001** | **0.0318** | **0.0130** | >0.9999 | 0.1355 | 0.4773 |
|  | OB vs V2 | 0.2539 | **<0.0001** | 0.9989 | 0.6463 | 0.0977 | **0.0106** | 0.4965 |
|  | M1 vs S1 | 0.8384 | **0.0001** | 0.9651 | 0.9854 | 0.1181 | 0.0787 | 0.9999 |
|  | M1 vs V2 | 0.5548 | **<0.0001** | **0.0072** | 0.8287 | **<0.0001** | **0.0051** | 0.9993 |
|  | S1 vs V2 | 0.9994 | 0.3830 | 0.0930 | 0.3875 | 0.1020 | 0.9561 | >0.9999 |

A. Statistical evaluation of the relative spectral power in function of the cortical region, behavioral state, and interaction between both factors. Repeated mixed-effects model. B. p values of the Sidak multiple comparisons test, comparing the differences in the relative power between the different cortical region of the right hemisphere. OB, olfactory bulb; M1, primary
motor cortex; S1, primary somato-sensory cortex; V2, secondary visual cortex; df, degrees of freedom; W, wakefulness; LG, low gamma; HG, high gamma; HFO, high frequency oscillations.

**Figure 4.**
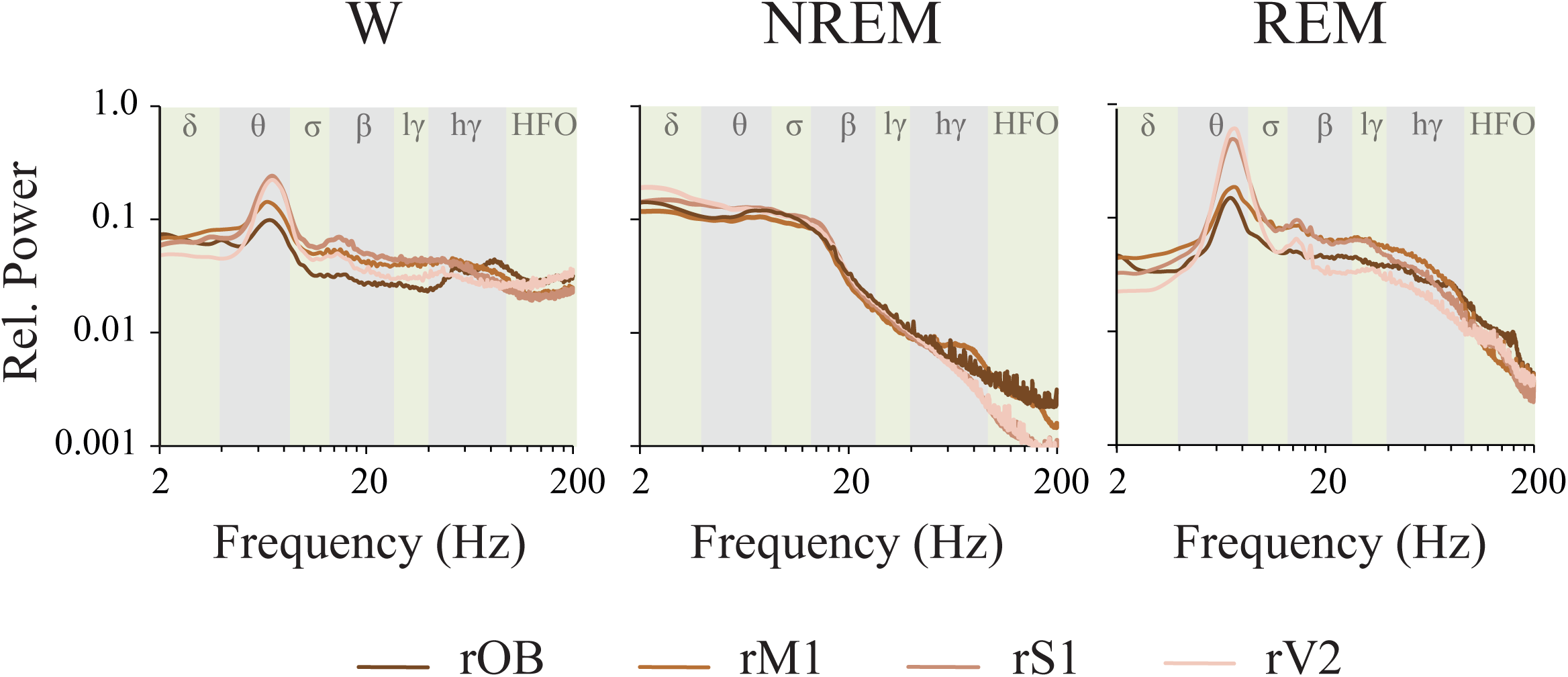
Relative power. Mean relative power profile of each behavioral state of the right hemisphere during the light period. This approach removes the effect of the distance between the active and the referential electrode. The analyzed frequency bands are indicated by different colors in the background of the graphics. OB, olfactory bulb; M1, primary motor cortex; S1, primary somato-sensory cortex; V2, secondary visual cortex; W, wakefulness; r, right; l, left; lγ, low gamma or LG; hγ, high gamma or HG; HFO, high frequency oscillations.

The most remarkable differences in the relative power between the brain regions were noticed during REM sleep, where V2 and S1 theta power was greater than in OB and M1. Another interesting finding was that LG and HG in M1 were higher than in V2. During NREM sleep, S1 power was higher than OB for the delta frequency band, and larger than OB and M1 for the theta band. On the other hand, the power in the OB was lower than M1 and S1 for LG, but became higher for HG and HFO during W.

### 2.2. Power spectrum: light Vs. dark phases

Next, we examined the effects of the light/dark phases on the iEEG oscillatory activity for the right hemisphere. Figure 5 shows the light/dark predominance (see the procedure in the Figure legend; the same approach was used to show the data in next figures). We can readily observe that the classic frequency bands show significant differences only during NREM sleep; beta and LG power were larger during the dark than during the light phase in M1. Employing a more precise evaluation using the empirical cluster analysis, we determined that during NREM sleep in M1, clusters of frequencies that include sigma, beta, LG, HG and HFO bands were larger in the dark phase. A cluster of frequencies within the HFO band also increased during the dark phase in the OB during this behavioral state. Regarding REM sleep, although no changes were found in the classical frequency bands, we found that a specific cluster within theta frequency band which was higher during the day than during the night in V2.

**Figure 5.**
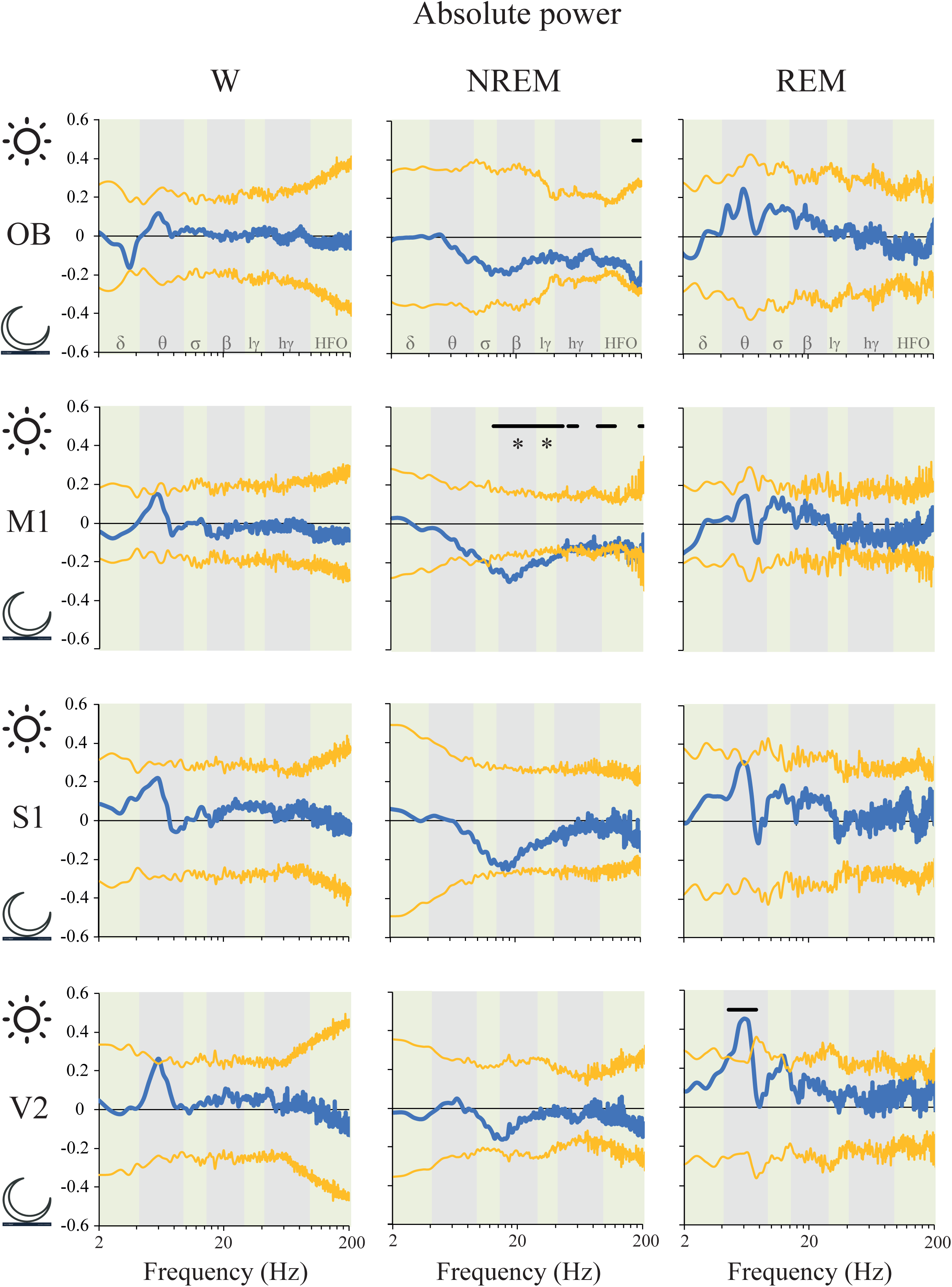
Absolute power: light Vs. dark phases differences. The predominance was calculated by means of the formula: (a-b)/(a+b). “a” represents the mean power for each frequency in the light phase, and “b” the mean power in the dark period. A positive value means that power during the light period was higher than during dark period and *vice versa*. The blue traces indicate the mean power difference between light and dark phases. The yellow lines represent the standard deviation of the mean with respect to zero. The statistical evaluation was performed by the two-tailed paired t-test with Bonferroni correction for multiple comparisons; * indicates significant differences, p < 0.0071. We also performed a data driven approach comparing empirical clusters of frequencies; black lines represent statistical differences in cluster of frequencies, p < 0.05. In M1 the following frequency clusters were significantly larger during NREM sleep in the dark phase: 13 to 46 Hz (p = 0.001), 51 to 60.5 Hz (p = 0.016) 87 to 92 Hz (p = 0.038), 93.5 to 100 Hz (p = 0.025), 101 to 111 Hz (p = 0.012), 112.5 119.5 Hz (p = 0.024),124 to 125.5 Hz (p = 0.047), 186.5 to 192 Hz (p = 0.036) and 193 to 200 Hz (p = 0.029). In the OB during NREM sleep, the power of the frequencies 171.5 to 187.5 and 188.5 to 199.5 Hz were larger during the night (p = 0.018 and p = 0.047, respectively). During REM sleep, the cluster 4.5 to 7.5 Hz, was higher during the day than during the night in V2 (p = 0.038). The analysis was performed for the right hemisphere. OB, olfactory bulb; M1, primary motor cortex; S1, primary somato-sensory cortex; V2, secondary visual cortex; lγ, low gamma or LG; hγ, high gamma or HG; HFO, high frequency oscillations.

### 2.3. Power spectrum: right Vs. left hemispheres

iEEG absolute power laterality was analyzed for both the light (Supplementary Figure 5) and dark (Supplementary Figure 6) phases. Neither t-test evaluation of classical frequency bands nor cluster analysis showed statistical differences between right and left hemispheres, either during W or sleep.

**Figure 6.**
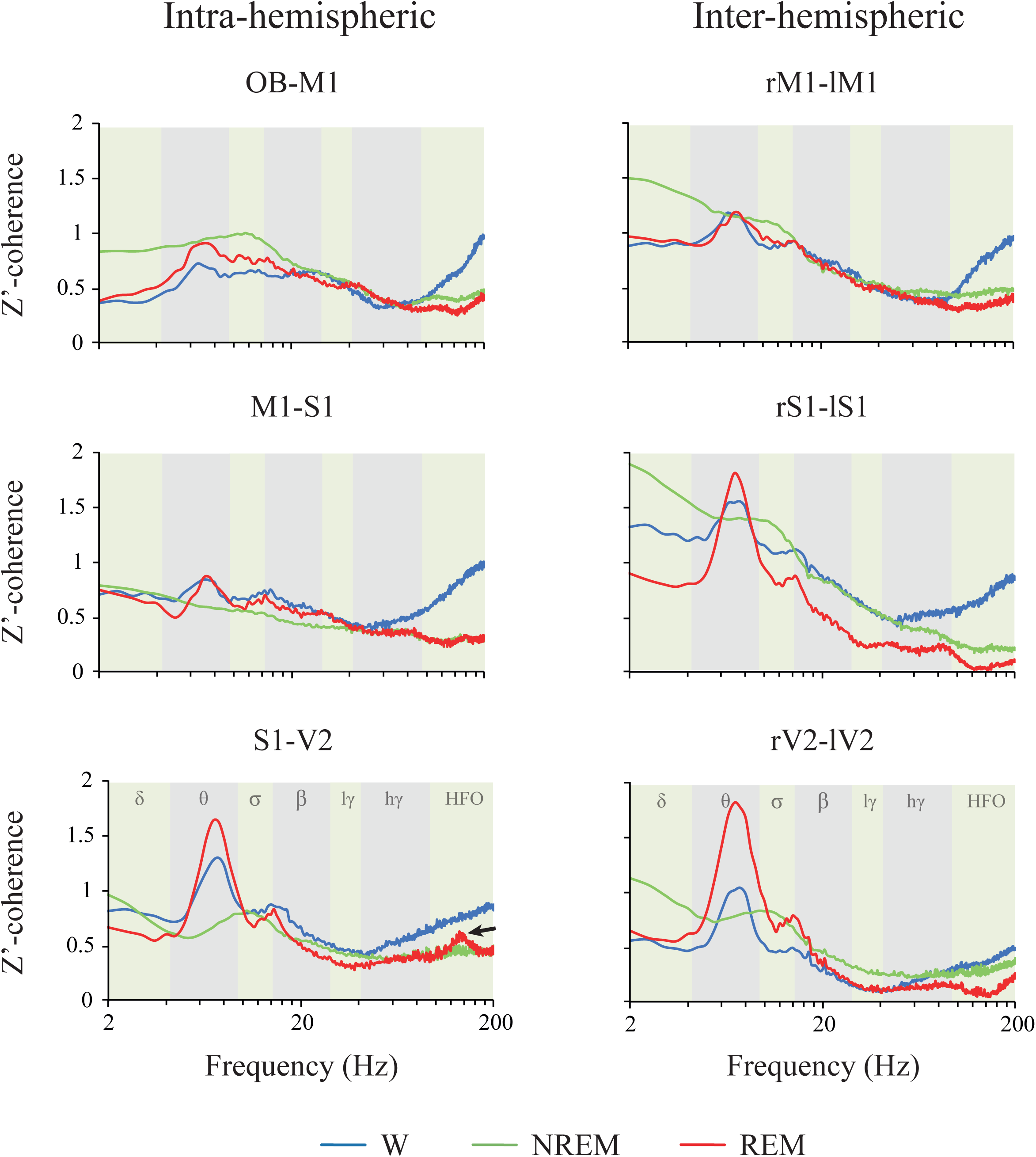
Z’-coherence. Mean z’-coherence profile of the inter-hemispheric and intra-hemispheric derivations (between adjacent areas) during wakefulness (W), NREM and REM sleep in the light phase. The analyzed frequency bands are indicated by different colors in the background of the graphics. OB, olfactory bulb; M1, primary motor cortex; S1, primary somato-sensory cortex; V2, secondary visual cortex; r, right; l, left; lγ, low gamma or LG; hγ, high gamma or HG; HFO, high frequency oscillations.

### 2.4. Coherence: effects of behavioral states and derivations

The z’-coherence during W, NREM and REM sleep for the right intra-hemispheric and the inter-hemispheric combination of adjacent electrodes during the light period is shown in Figure 6. In Supplementary Figure 7, the data were re-plotted in order to appreciate the differences in function of the derivations. The statistical results of the repeated measures mixed-effects model are shown in Table 3. Interestingly, no significant differences in z’-coherence were observed in function of the derivation, except for delta and theta bands. However, behavioral states and the interaction between localization and behavioral states modified z’-coherence in all the frequency bands (Table 3).

**Table 3.** Z’-coherence.

| Frequency | Derivation |  |  | Behavioral State |  |  | Derivation x State |  |  |
| --- | --- | --- | --- | --- | --- | --- | --- | --- | --- |
|  | df | p | F | df | p | F | df | p | F |
| Delta | 5,50 | **0.0411** | 2.5 | 2, 20 | **<0.0001** | 23.1 | 10, 82 | **0.0121** | 2.5 |
| Theta | 5,50 | **0.0423** | 2.5 | 2, 20 | **0.0015** | 9.2 | 10, 82 | **<0.0001** | 10.7 |
| Sigma | 5,50 | 0.1992 | 1.5 | 2, 20 | **0.0221** | 4.6 | 10, 82 | **0.0018** | 3.2 |
| Beta | 5,50 | 0.3609 | 1.86 | 2, 20 | **0.0356** | 3.9 | 10, 82 | **0.0129** | 2.4 |
| LG | 5,50 | 0.3687 | 1.1 | 2, 20 | **0.0058** | 6.7 | 10, 82 | **0.0094** | 2.6 |
| HG | 5,50 | 0.2258 | 1.4 | 2, 20 | **0.0004** | 11.7 | 10, 82 | **<0.0001** | 4.3 |
| HFO | 5,50 | 0.0600 | 2.3 | 2, 20 | **<0.0001** | 32.1 | 10, 82 | **<0.0001** | 5.5 |

Statistical evaluation of the z'-coherence in function of the derivation, behavioral state, and interaction between both factors. Repeated measures mixed-effects model. df, degrees of freedom; LG, low gamma; HG, high gamma; HFO, high frequency oscillations.

**Figure 7.**
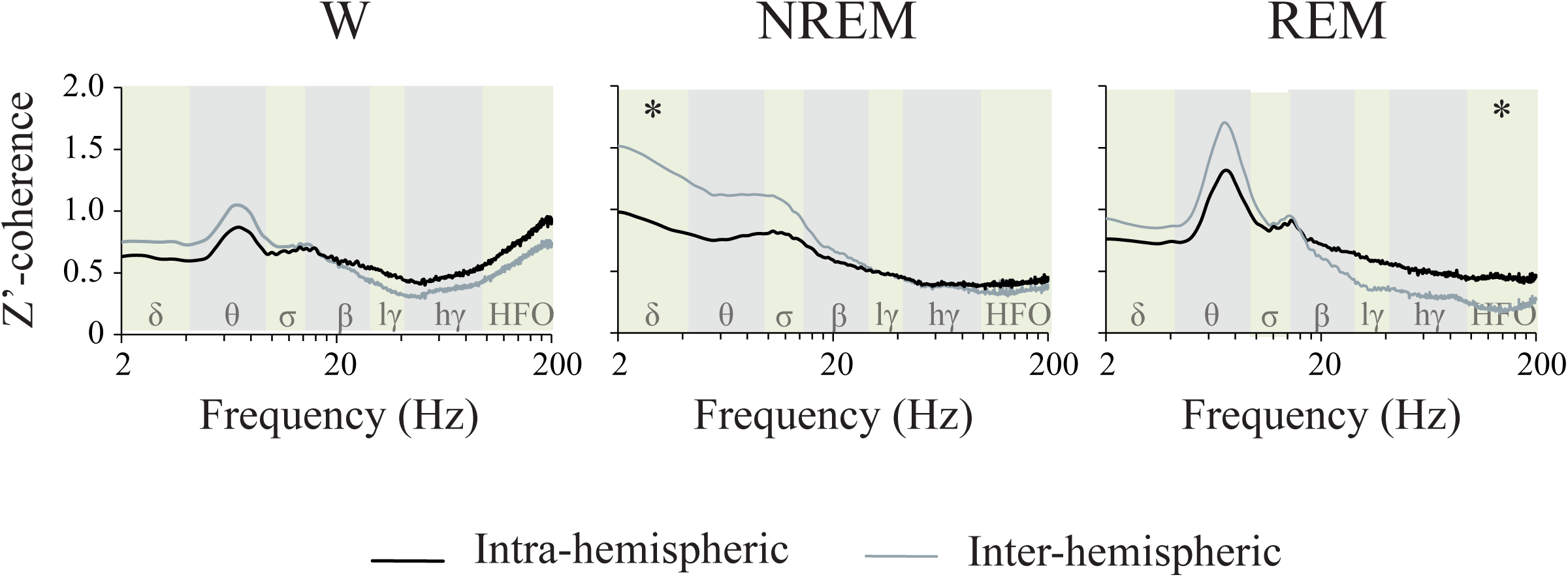
Z’-coherence differences between intra- and inter-hemispheric derivations. z’-coherence profile of the mean intra-hemispheric (OB-M1, M1-S1 and S1-V2) and inter-hemispheric (right-left M1, S1, V2) derivations during wakefulness (W), NREM and REM sleep in the light phase. The analyzed frequency bands are indicated by different colors in the background of the graphics. OB, olfactory bulb; M1, primary motor cortex; S1, primary somato-sensory cortex; V2, secondary visual cortex; r, right; l, left; lγ, low gamma or LG; hγ, high gamma or HG; HFO, high frequency oscillations. Asterisks indicate significant differences, p < 0.05. M1, primary motor cortex; S1, primary somato-sensory cortex; V2, secondary visual cortex; lγ, low gamma or LG; hγ, high gamma or HG; HFO, high frequency oscillations.

The spectral z’-coherence of each frequency band differences between W, NREM and REM sleep for each electrode combination is also shown in Figure 3; the p values for each comparison are shown in Supplementary Figure 8. The most important results are the following. Delta z’-coherence during NREM sleep was larger than during the other behavioral states in most derivations. Moreover, sigma z’-coherence had higher values during NREM than W in the intra-hemispheric combination OB-M1 and the inter-hemispheric sensory cortices. Theta z’-coherence increased during REM sleep in comparison to NREM sleep in the posterior intra and inter-hemispheric combination of electrodes. HG and HFO intra and inter-hemispheric z’-coherence was larger during W compared to REM and NREM sleep in most derivations. Also, visual inter-hemispheric LG and HG z’-coherence were lower during REM sleep compared to NREM sleep.

**Figure 8.**
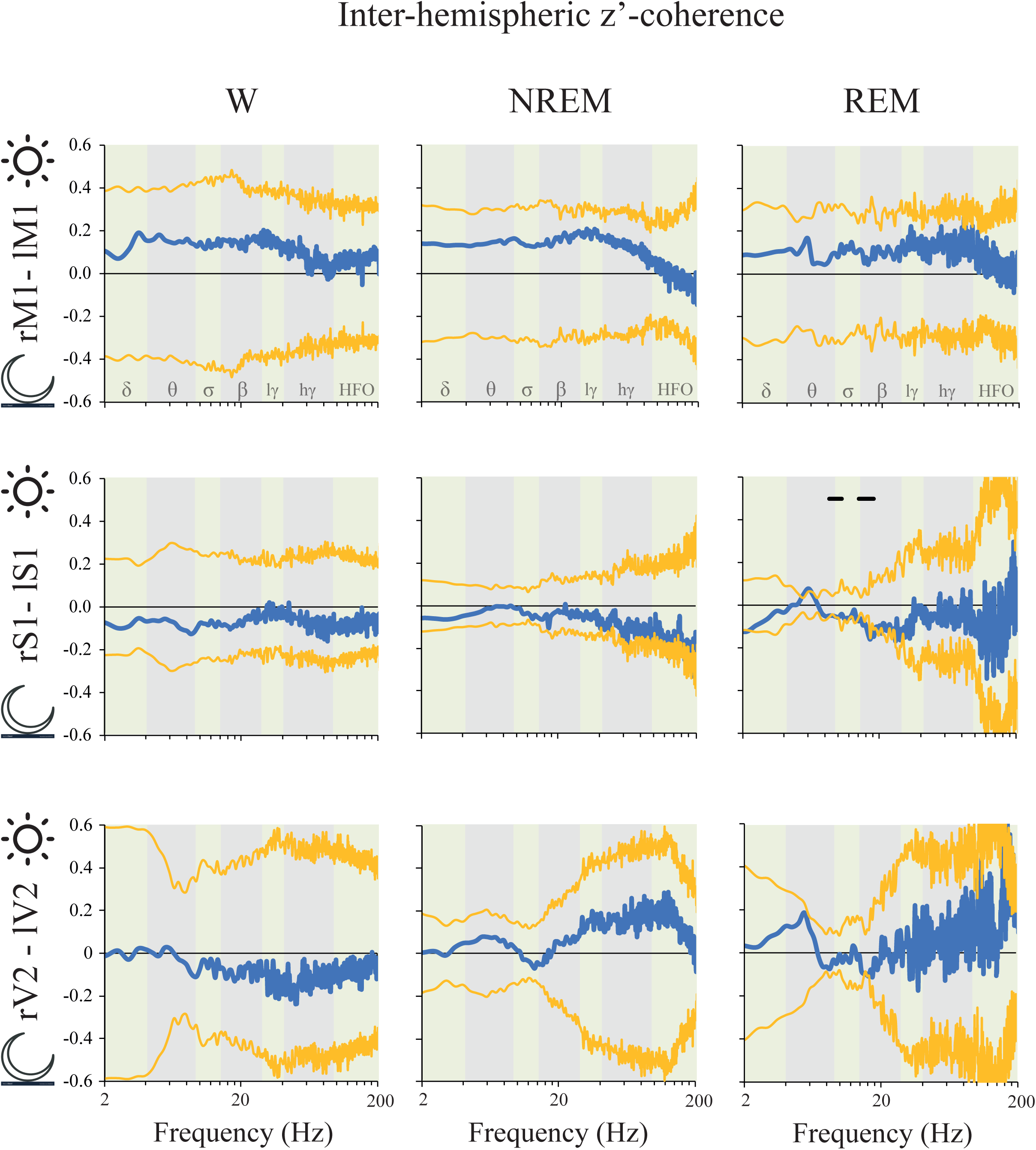
Inter-hemispheric z’-coherence: light Vs. dark phases differences. The predominance was calculated by means of the formula: (a-b)/(a+b). “a” represents the mean z’-coherence for each frequency in the light phase, and “b” the mean z’-coherence in the dark period. A positive value means that z’-coherence in light period was higher than during dark period and *vice versa*. The blue traces indicate the mean coherence difference between light and dark phases. The yellow lines represent the standard deviation of the mean with respect to zero. The statistical evaluation was performed by the two-tailed paired t-test with Bonferroni correction for multiple comparisons; no significant differences were observed. We also performed a data driven approach comparing empirical clusters of frequencies; black lines represent statistical differences in cluster of frequencies, p < 0.05. During REM sleep, inter-hemispheric S1 z’-coherence was higher during the dark phase for the clusters 8.5 to 10.5 Hz (p = 0.008) and 14 to 18.5 Hz (p = 0.005). M1, primary motor cortex; S1, primary somato-sensory cortex; V2, secondary visual cortex; r, right; l, left; lγ, low gamma or LG; hγ, high gamma or HG; HFO, high frequency oscillations.

When comparing z’-coherence in function of the derivation, we could detect very few differences during W and sleep, and in most of the cases included differences between intra Vs. inter-hemispheric derivations (the complete statistical analysis is shown in Supplementary Figure 9). Therefore, we analyzed the differences between the average inter-hemispheric (right and left M1, S1 and V2 derivations) and intra-hemispheric z’-coherences (OB-M1, M1-S1 and S1-V2 derivations) during W and sleep (Figure 7). We found a significant effect of the interaction between the behavioral state and the derivation type for delta (F _(2,14)_ = 4.7, p = 0.028) and HFO bands (F _(2,14)_ = 3.8, p = 0.048). During NREM sleep delta inter-hemispheric was higher than the intra-hemispheric z’-coherence (p = 0.046). In contrast, during REM sleep HFO intra-hemispheric was larger than the inter-hemispheric z’-coherence (p = 0.038).

**Figure 9.**
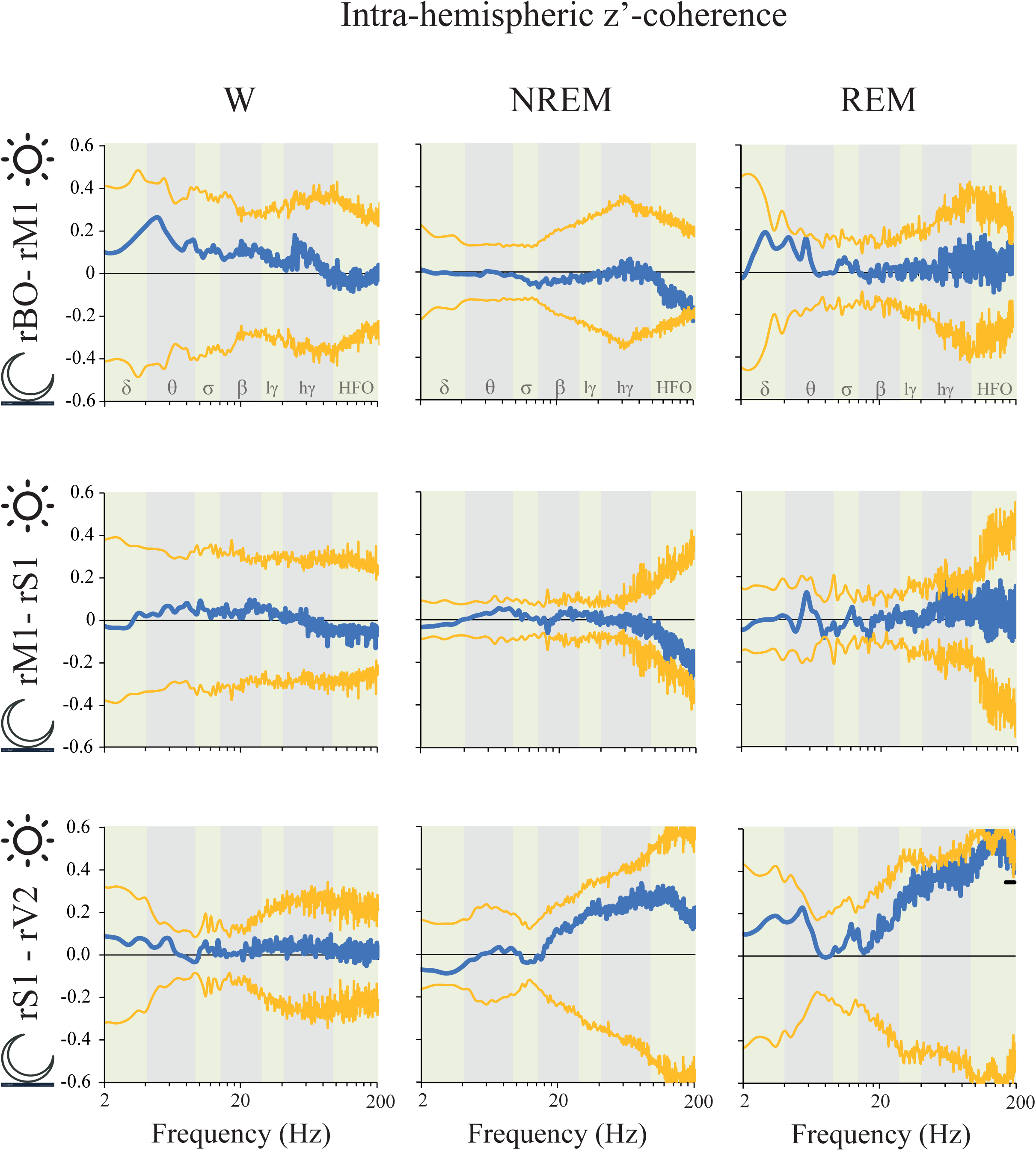
Intra-hemispheric z’-coherence: light Vs. dark phases differences. The predominance was calculated by means of the formula: (a-b)/(a+b). “a” represents the mean z’-coherence for each frequency in the light phase, and “b” the mean z’-coherence in the dark period. A positive value means that z’-coherence in light period was higher than during dark period and *vice versa*. The blue traces indicate the mean z’-coherence difference between light and dark phases. The yellow lines represent the standard deviation of the mean with respect to zero. The statistical evaluation was performed by the two-tailed paired t-test with Bonferroni correction for multiple comparisons; no significant differences were observed. We also performed a data driven approach comparing empirical clusters of frequencies; black lines represent statistical differences in cluster of frequencies, p < 0.05. During REM sleep, intra-hemispheric S1-V2 z’-coherence was higher during the light phase for the cluster 173-200 Hz (p = 0.005). OB, olfactory bulb; M1, primary motor cortex; S1, primary somato-sensory cortex; V2, secondary visual cortex; r, right; lγ, low gamma or LG; hγ, high gamma or HG; HFO, high frequency oscillations.

### 2.5. Coherence: light Vs. dark phases

The influence of light/dark phases on z’-coherence was also analyzed. There were no significant differences for the classical frequency bands either for inter (Figure 8) or intra-hemispheric (Figure 9) z’-coherence between the dark and light phases during either W or NREM sleep. However, during REM sleep inter-hemispheric S1 z’-coherence was higher in the dark phase for two clusters of frequencies: 8.5 to 10.5 Hz and 14 to 18.5 Hz (Figure 8). Moreover, REM sleep intra-hemispheric S1-V2 z’-coherence was higher during the light phase for the cluster 173-200 Hz (Figure 9).

### 2.6. Coherence: right Vs. left hemispheres

Neither t-test analysis of classical bands nor cluster analysis showed intra-hemispheric z’-coherence differences between right and left hemispheres during the light phase (Supplementary Figure 11).

During the dark phase, a cluster circumscribed mainly within the theta band in S1-V2 derivation was higher in the right hemisphere during REM sleep. In contrast, clusters of frequencies within LG, HG and HFO were higher in the left hemisphere (Figure 10).

**Figure 10.**
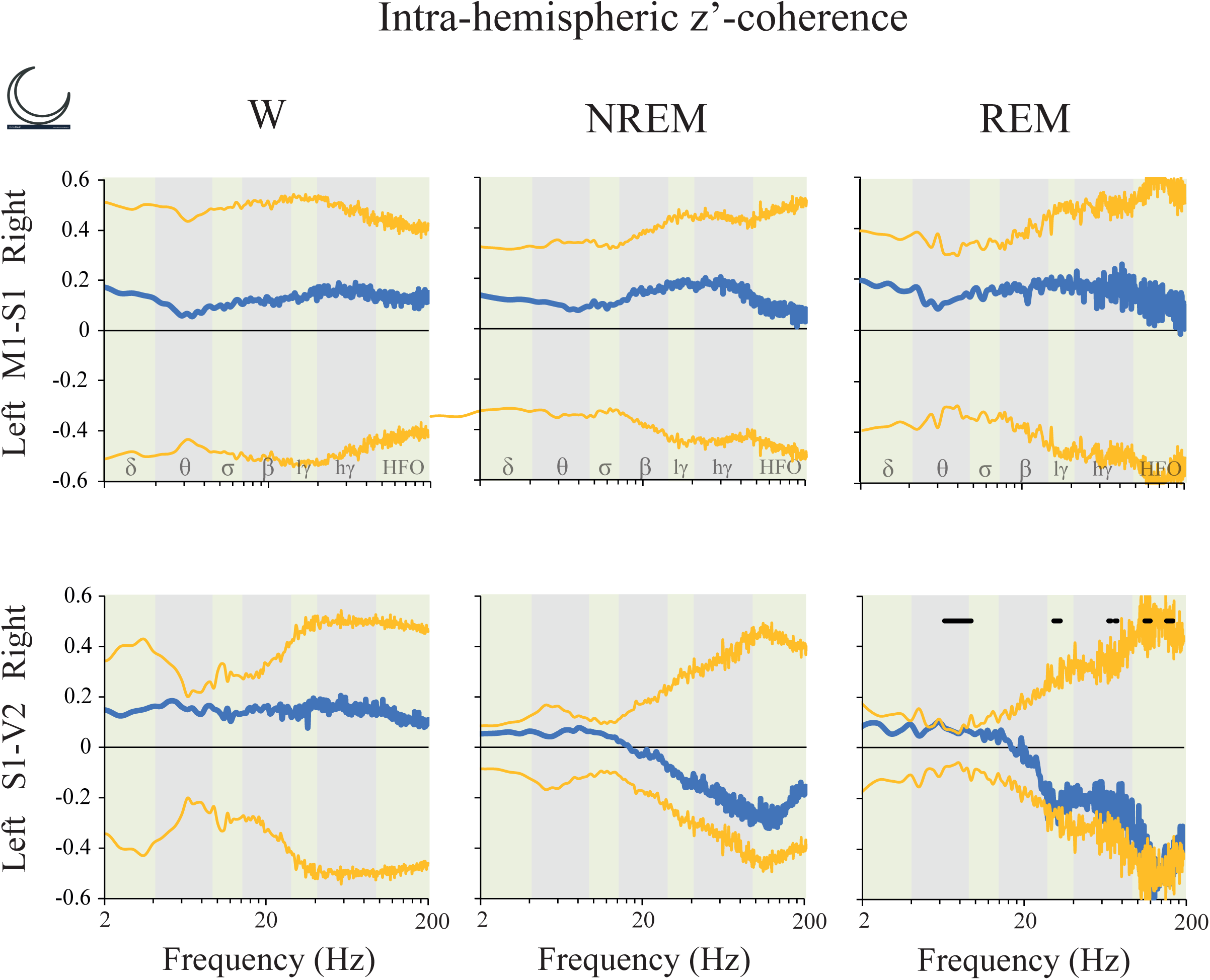
Intra-hemispheric z’-coherence: right Vs. left hemispheric difference during the dark phase. The predominance was calculated by means of the formula: (a-b)/(a+b). “a” represents the mean z’-coherence for each frequency for the right hemisphere, and “b” the mean z’-coherence of the left hemisphere. A positive value means that z’-coherence during light period was higher than during dark period and *vice versa*. The blue traces indicate the mean z’-coherence difference between light and dark phases. The yellow lines represent the standard deviation of the mean with respect to zero. The statistical evaluation was performed by the two-tailed paired t-test with Bonferroni correction for multiple comparisons; no significant differences were observed. We also performed a data driven approach comparing empirical clusters of frequencies; black lines represent statistical differences in cluster of frequencies, p<0.05. Coherence was higher in the right hemisphere for the cluster between 6.5 to 9.5 Hz (p= 0.0009) during REM sleep. In contrast, the clusters between 31 to 34 Hz (p = 0.022), 68 to 70 Hz (p = 0.042), 74.5 to 76.5 Hz (p = 0.048), 114 to 117 Hz (p = 0 .027), 118 to 123 Hz (p = 0.015), 124 to 151.5 (p = 0.001), 155.5 to 164 (p = 0.008) and 165 to 170 Hz (p = 0.017) were higher in the left hemisphere for the same behavioral state. M1, primary motor cortex; S1, primary somato-sensory cortex; V2, secondary visual cortex; lγ, low gamma or LG; hγ, high gamma or HG; HFO, high frequency oscillations.

## 3. Discussion

In the present study, we performed an exhaustive analysis of the power and coherence of the iEEG signal of the rat, as well as the impact of behavioral states, cortical areas, laterality (differences between hemispheres) and light/dark phases. iEEG power and coherence were largely affected by behavioral states and recording sites. On the contrary, the influence of the light/dark phases was detected only during sleep. Finally, while we did not find right/left differences in power either in W or sleep, we observed that intra and inter-hemispheric coherence differs between both hemispheres during REM sleep.

### 3.1. Technical considerations

We performed monopolar recordings (referenced to the cerebellum) utilizing screws (1 mm diameter) in contact with the dura mater as recording electrodes. With this recording design, we observed an important impact of the cortical site. However, as mentioned in Results section, the absolute power is highly dependent on the distance between the recording and reference electrodes. In order to discern the relative weight of the power of specific frequency bands in each channel, we also computed the relative power (absolute individual frequencies power normalized by total power). Hence, although complete analyses are provided in Supplementary Data, in the description of the result we did not focus on the absolute power as a function of the cortical area or in the total power. In other words, we emphasized the absolute power in function of behavioral states, and the relative power in function of the electrode site.

However, it is important to note that total power in OB is lower than in M1 and S1 during NREM sleep (Supplementary Figure 3); this result is not an artifact of the distance between the active and referential electrodes (that is the longest in this case), indicating that the amplitude of the slow waves during NREM sleep in the paleocortex is lower than in the neocortex.

The present as well as most of the studies that analyzed the iEEG spectrum focus on “classical” or “standard” frequency bands, that are associated with behavioral states and cognitive functions (Castro et al., 2013; Castro et al., 2014; Cavelli et al., 2017; Cavelli et al., 2018; Mahjoory et al., 2019; Nyhus and Curran, 2010). Nevertheless, the effect of different variables on power and coherence could be circumscribed to only a subset of frequencies within a “classical” band or could include changes that extend over the classical bands limits. In these cases, the results could be washed out when the whole band is analyzed (Myrden and Chau, 2016). Because of this, for day/night and laterality analyses we also performed the result-driven analysis of clusters of frequencies. This methodology allowed us to unveil changes in the iEEG power that not coincide exactly with “standard” or “classical” frequency bands.

### 3.2. iEEG power

During W, gamma and HFO power reached the maximal level (Cavelli et al., 2015; Cavelli et al., 2018); however, although in our analysis we eliminated the epochs with movements artifacts, we can’t rule out the possibility that part of the signal corresponds to muscle activity contamination that reaches cortical electrodes through volume conduction. Interestingly, in the OB two small deflections in the HG band during W (signaled by blue arrows in Figure 2A) are readily observed. Gamma oscillations in the OB are known since the pioneering study of (Adrian, 1942); these oscillations are in phase with the respiratory potentials and significantly increase during active exploration (Cavelli et al., 2019; Zhuang et al., 2019). In accordance with this, we found that HG power was higher in the OB during W than during sleep.

NREM sleep was characterized by a higher power spectrum profile associated with large absolute and relative power values in slow frequency bands (delta, theta and sigma), that can be observed throughout the cortex (Figure 2A and B). A remarkable change in the slope is readily observed at ≈ 16 Hz; from this point, there is a marked constant decrease in power as a function of the frequency. As mentioned in the Introduction, delta (and also low theta) is related to the cortical and thalamic slow oscillations, while sigma power to sleep spindles; both electrographic features that characterize NREM sleep. As expected, delta and sigma band power during NREM were larger than during W and REM sleep in most of the cortical areas. Furthermore, we found that absolute theta power was higher in NREM than during W and REM sleep in the motor and somatosensory cortices. Hence, even if in W and REM sleep exhibit a clear peak at ≈ 7 Hz in posterior areas, and the relative theta rhythm predominates over other frequency bands, the absolute theta power (considering the whole band) is not as high as in NREM sleep.

During REM sleep, the relative weight of the theta band is highlighted with the analysis of the relative power (Figure 4). The main origin of theta rhythm is in the hippocampus; this hippocampal theta rhythm modulates cortical neuronal activity (Cavelli et al., 2018; Gonzalez et al., 2020b; Sirota et al., 2008; Winson, 1974). A prominent peak in theta and relatively large power in gamma and HFO (larger than during NREM sleep) characterize REM sleep (Cavelli et al., 2018). In fact, in the present study we found that HG power was higher during REM than during NREM sleep in the motor cortex. Also, as described before (Cavelli et al., 2018), a narrow peak in HFO at ≈ 130 Hz can be appreciated during REM sleep in the OB and sensory cortices (signaled with red arrows in Figure 2). HFO is implicated in the sensory processing (Bauer et al., 2014), and a recent study suggest that the OB is a source of HFO (Hunt et al., 2019). However, the whole HFO band power was not significantly different between NREM and REM sleep, probably because the set of frequencies involved in the peak is much narrower than the whole band.

### 3.3. iEEG coherence

The spectral coherence is a tool to examine the functional interactions between different cortices as a function of the frequency (Bullock and McClune, 1989; Castro et al., 2013). In accordance with previous studies (Castro et al., 2013; Cavelli et al., 2015; Cavelli et al., 2018), during W large values of intra and inter-hemispheric coherence were observed for high frequencies (HG and HFO) in almost all the derivations. Hence, during W, both gamma and HFO power (that reflects local synchronization) and coherence (that suggest synchronization between areas) are high.

During NREM sleep, delta coherence was higher than during W and REM sleep; thus, large delta power and coherence characterize NREM sleep. However, (Pal et al., 2016) found that during NREM there was a reduction in the cortico-cortical delta coherence in comparison to W. Of note is that these authors used the mean global coherence (an average of the coherence for the individual channel pairs), and they only evaluated inter-hemispheric combination of electrodes. It was interesting the fact that delta coherence was larger in homologous inter-hemispheric than in intra-hemispheric derivations (Figure 7); this fact was also demonstrated in humans (Achermann and Borbely, 1998).

Similar to our previous report (Cavelli et al., 2018), we found that REM sleep is characterized by large theta coherence, especially between visual and/or somatosensory electrodes. Another valuable issue is that HG and HFO coherence during REM sleep is lower than in W, as was previously described in rats (Cavelli et al., 2015; Pal et al., 2016). Also in cats, there are very low gamma coherence values (both LG and HG) during REM sleep (Castro et al., 2013; Castro et al., 2014). In accordance with (Cavelli et al., 2018), HFO coherence for intra-hemispheric posterior (S1-V2, sensory) derivations has a clear “peak” during REM sleep (indicated by a black arrow in Figure 6); this peak did not reach statistical significance in comparison to NREM sleep when we analyzed the whole HFO band. In spite of this, it is important to note that HFO intra-hemispheric z’-coherence was significantly higher than inter-hemispheric coherence during REM sleep (Figure 7).

### 3.4. Impact of the light/dark phases

In the present report, we analyzed the iEEG during the subjective day (9 A.M. to 3 P.M) and compared it with the subjective night (9 P.M to 3 A.M.); the lights were on from 6 AM to 6 PM. Hence, we evaluated the average of 6 h periods of the light/dark phases in the middle of these phases. In other words, the 3 h at the beginning and at the end of the phases, that should be more unsteady, were not analyzed. This is important to take it into account because previous studies showed important modification in the hour to hour iEEG oscillations (Osorio et al., 2020; Vyazovskiy et al., 2002).

Albino rats have short sleep cycles (on average ≈ 11 minutes) and are more active during the dark phase; i.e., light phase is their main resting period (Datta and Hobson, 2000; Stephenson et al., 2012; Trachsel et al., 1991). Interestingly, during REM sleep there was a clear predominance of theta power (≈ 5-7 Hz) in the light phase (Figure 5); this effect reached the maximum in V2, and it was statistically significant between 4.5 to 7.5 Hz. Theta amplitude and frequency variates during REM sleep (Karashima et al., 2004), and this power predominance during the light phase could be explained by a deeper REM sleep in the resting phase. In contrast, during NREM sleep high frequencies power of the iEEG were skewed toward the dark phase (Figure 5). These phenomena could be related to a shallow NREM sleep during the active phase.

To the best of our knowledge, the light/dark phases differences in the iEEG spectral coherence was not studied before. Cluster analysis revealed that there were light/dark phases differences, but only during REM sleep. We found a dark phase predominance within theta, sigma and beta bands in S1 inter-hemispheric derivation (Figure 7). In contrast, there was a light phase predominance for frequencies higher than 173 Hz in S1-V2 intra-hemispheric derivation during this behavioral state (Figure 8). The functional meaning of this day/night differences in electro-cortical coherence confined just to REM sleep is unknown, but may be related to the circadian strength of the cognitive function executed during this behavioral state, such as memory processing (Poe, 2017; Sara, 2017).

### 3.5. Right/left hemispheric differences in electro-cortical activity

There were no differences between the right/left hemispheres iEEG power either during light or dark phases during W. Vyazovskiy and Tobler (2008) described iEEG laterality at 4.5-6.0 Hz during a hand preference task; however, as is in our results, the authors did not find differences in naïve animals.

During NREM sleep, Vyazovskiy et al. (2002) showed a left-hemispheric predominance of low-frequency power in the parietal cortex at the beginning of the light period, when sleep pressure is high. The left-hemispheric dominance changed to a right-hemispheric dominance in the course of the resting phase, when sleep pressure dissipated. Also, during recovery from sleep deprivation, parietal left-hemispheric predominance was enhanced. We did not see hemispheric laterality in any frequency band, probably because we did not record the first 3 hours of the light phase, that seems to be the time where laterality is mostly developed.

During REM sleep, right-hemispheric predominance in the theta band power has been described (Vyazovskiy et al., 2002). Although, no significant differences were observed, in accordance with these authors, there was a clear tendency for right predominance in the theta band power in M1 and S1 during REM sleep, both during light and dark phases (Supplementary Figures 5 and 6).

We did not find previous reports that compared the effect of laterality in the iEEG coherence in rodents; however, changes in iEEG coherence during W have been shown in humans in relation to their skilled hand (Boldyreva and Zhavoronkova, 1991). In the present report, right/left significant differences in coherence were found only during REM sleep in the dark phase (Figure 10). We found a right predominance of frequencies within the theta range in S1-V2 derivation, and a left predominance for clusters within gamma and HFO bands. New experimental approaches are needed to explain these differences between both hemispheres.

In conclusion, in the present study, we carried out a thorough analysis of the spectral power and coherence of the rat iEEG. We found major effects on these parameters in function of both behavioral states and cortical areas. We also revealed that there are night/day differences in power and coherence during sleep, but not during W. Additionally, while we did not find right/left differences in power either in W or sleep, we observed that during REM sleep intra-hemispheric coherence differs between both hemispheres. We consider that this systematic analysis of the iEEG dynamics during physiological W and sleep provides a template or reference for comparison with pharmacological, toxicological or pathological challenges.

## 4. Material and Methods

### 4.1. Experimental Animals

Eleven Wistar male adult rats (270-300 g) were used for this study. The animals were determined to be in good health by veterinarians of the institution. All experimental procedures were conducted in agreement with the National Animal Care Law (#18611) and with the “Guide to the care and use of laboratory animals” (8th edition, National Academy Press, Washington DC., 2010). Furthermore, the Institutional Animal Care Committee approved the experimental procedures (No 070153-000332-16). Adequate measures were taken to minimize pain, discomfort or stress of the animals, and all efforts were made to use the minimal number of animals necessary to obtain reliable scientific data. Animals were maintained on a 12-h light/dark cycle (lights on at 6.00 A.M.) and housed five to six per cage before the experimental procedures. Food and water were freely available.

### 4.2. Surgical Procedures

We employed surgical procedures similar to those used in our previous studies (Cavelli et al., 2018; Gonzalez et al., 2019; Mondino et al., 2019). The animals were chronically implanted with intracranial electrodes. Anesthesia was induced with a mixture of ketamine-xylazine (90 mg/kg; 5 mg/kg i.p., respectively). Rats were positioned in a stereotaxic frame and the skull was exposed. In order to record the iEEG, stainless steel screw electrodes were placed on the skull above the right and left M1, S1 and V2, as well as on the right OB and cerebellum (reference electrode). A representation of the electrodes’ position and their coordinates according to (Paxinos and Watson, 2005), are shown in Figure 1A and B. In order to record the electromyogram (EMG), a bipolar electrode was inserted into the neck muscle. The electrodes were soldered into a 12-pin socket and fixed to the skull with acrylic cement. At the end of the surgical procedures, an analgesic (Ketoprofen, 1 mg/kg subcutaneous) was administered. Incision margins were kept clean and a topical antibiotic was applied on a daily basis. After the animals recovered from the preceding surgical procedures, they were adapted to the recording chamber for one week.

### 4.3. Sleep recordings

Animals were housed individually in transparent cages (40 × 30 x 20 cm) containing wood shaving material in a temperature-controlled (21-24 °C) room, with water and food *ad libitum*, under a 12:12 hs light/dark cycle (lights on at 6 A.M.). Experimental sessions were conducted during the light (9 A.M. to 3 P.M.) and dark periods (9 P.M. to 3 A.M.) in a sound-attenuated chamber that also acts as a Faraday box. The recordings were performed through a rotating connector, to allow the rats to move freely within the recording box. Bioelectric signals were amplified (×1000), filtered (0.1-500 Hz), sampled (1024 Hz, 16 bits) and stored in a PC using Spike 2 software (Cambridge Electronic Design).

### 4.4. Data analysis

The states of sleep and W were determined in 10 s epochs. W was defined as low voltage fast waves in frontal cortex, a mixed theta rhythm in occipital cortex and relatively high EMG activity. Light sleep (LS) was determined as high voltage slow cortical waves interrupted by low voltage fast iEEG activity. Slow wave sleep (SWS) was defined as continuous high amplitude slow (1-4 Hz) neocortical waves and sleep spindles combined with a reduced EMG activity. LS and SWS were grouped as NREM sleep. REM sleep was defined as low voltage fast frontal waves, a regular theta rhythm in parietal and occipital cortices, and a silent EMG except for occasional twitching. In order to analyze power spectrum (in each channel) and coherence (between pairs of iEEG channels or derivations) we used procedures similar to those done in our previous studies (Cavelli et al., 2015; Cavelli et al., 2018; Mondino et al., 2019). The maximum number of non-transitional and artifact-free periods of 30 seconds was selected during each behavioral state to determine the mean power and coherence for each rat. Power spectrum was estimated by means of the *pwelch* built-in function in MATLAB using the following parameters: window = 30s, noverlap = [], nfft = 2048, fs =1024, which correspond to employing 30s sliding windows with half window overlap with a 0.5 Hz resolution.

The coherence between selected pairs of iEEG channels was analyzed in 30 s epochs. We chose for the analysis all the intra-hemispheric and inter-hemispheric pairwise combination of adjacent cortices (the distance between adjacent neocortical electrodes was 5 mm; Figure 1B). For each period, the Magnitude Squared Coherence for each channel (for details about coherence definition see (Bullock and McClune, 1989; Castro et al., 2013), was calculated with the *mscohere* built-in MATLAB (parameters: window = 30s, noverlap, nfft = 2048, fs =1024). In order to normalize the data and conduct parametric statistical tests, we applied the Fisher z’ transform to the coherence values (Castro et al., 2013).

The analysis of the data was performed for the classically defined frequency band in rodents: delta, 1-4 Hz; theta, 5-9 Hz; sigma, 10-15 Hz; beta, 16-30 Hz; low gamma (LG), 31-48 Hz; high gamma (HG) 52-95 Hz; and high frequency oscillations (HFO), 105-200 Hz (Cavelli et al., 2018; Mondino et al., 2019). Frequencies around 50 and 100 Hz were not analyzed to avoid alternating current artifacts. Differences in mean power and coherence among states (W, NREM and REM sleep) and electrode position or derivation, were evaluated by means of a two-ways repeated measures mixed-effects model, and Sidak as a correction for multiple comparisons test. We employed a mixed-effects model because we had to remove the information of noisy iEEG channels in 3 rats. We also computed the relative power as the absolute power of a specific frequency band/ the sum of the power from 0.5 to 200 Hz. Statistical significance was set at p < 0.05.

In order to determine if power and coherence were different between the time of the day (light Vs dark phases) or between hemispheres (right Vs. left), a paired two-tailed Student test was performed for each of the abovementioned bands. As we analyzed seven frequency bands, a Bonferroni correction for multiple comparisons was applied. With this correction p<□0.0071 was considered statistically significant.

We are aware that the specific start- and end-points of each frequency band is arbitrary, and vary between subjects (Haegens et al., 2014; Myrden and Chau, 2016). Because of this, for the day/night and laterality analyses we also performed a second evaluation by means of a cluster-based permutation test, consisting of comparing empirical clusters of frequencies against a randomized distribution, thus allowing the frequency bands to be delimited in a statistical approach without the need of a previous convention (Gonzalez et al., 2020a).

## 5. Acknowledgements

This study was supported by grants from the Agencia Nacional de Investigación e Innovación (ANII) FCE-1-2011-1-5997, Comisión Sectorial de Investigación Científica (CSIC) I+D-2016-589, and Programa de Desarrollo de Ciencias Básicas (PEDECIBA), Uruguay.

## Abbreviations

EEG: electroencephalogram
iEEG: intra-cranial electroencephalogram
HFO: high frequency oscillations
HG: high gamma
LG: low gamma
M1: primary motor cortex
NREM: non-REM
OB: Olfactory bulb
REM: Rapid eye movement
S1: primary somatosensory cortex
V2: secondary visual cortex
W: Wakefulness

