## Supplementary material for "Power and coherence in the EEG of the rat: impact of behavioral states, cortical area, lateralization and light/dark phases"

A

| Cortex | Comparison | Delta | Theta | Sigma | Beta | LG | HG | HFO |
|---|---|---|---|---|---|---|---|---|
| **OB** | W vs NREM | 0.6445 | 0.4622 | **0.0123** | 0.9732 | **0.0199** | **<0.0001** | **<0.0001** |
| | W vs REM | 0.1889 | 0.7319 | 0.9566 | 0.4866 | 0.0957 | **<0.0001** | **<0.0001** |
| | NREM vs REM | **0.0138** | 0.0774 | **0.0031** | 0.2634 | 0.9026 | 0.8339 | 0.9684 |
| **M1** | W vs NREM | **<0.0001** | **0.0004** | **<0.0001** | 0.2875 | **0.0006** | **<0.0001** | **<0.0001** |
| | W vs REM | 0.6163 | 0.6545 | >0.9999 | 0.7885 | 0.1984 | **<0.0001** | **<0.0001** |
| | NREM vs REM | **<0.0001** | **<0.0001** | **<0.0001** | **0.0464** | 0.1231 | **0.0133** | 0.3058 |
| **S1** | W vs NREM | **<0.0001** | **0.0064** | **<0.0001** | 0.1379 | **0.0321** | **<0.0001** | **<0.0001** |
| | W vs REM | 0.8051 | 0.8555 | 0.9998 | 0.9115 | 0.6745 | **0.0020** | **<0.0001** |
| | NREM vs REM | **<0.0001** | **0.0474** | **<0.0001** | **0.0331** | 0.3065 | 0.0716 | 0.9649 |
| **V2** | W vs NREM | 0.1540 | 0.8113 | 0.1495 | 0.9978 | 0.1898 | **0.0023** | **0.0002** |
| | W vs REM | 0.9614 | 0.7019 | 0.9994 | 0.8749 | 0.1905 | **0.0451** | **0.0004** |
| | NREM vs REM | 0.0562 | 0.9973 | 0.1182 | 0.9398 | >0.9999 | 0.6695 | 0.9937 |

B

| States | Comparison | Delta | Theta | Sigma | Beta | LG | HG | HFO |
|---|---|---|---|---|---|---|---|---|
| **W** | OB vs M1 | 0.9741 | 0.9921 | 0.9767 | 0.5754 | 0.2381 | 0.0723 | **0.0001** |
| | OB vs S1 | 0.8577 | 0.2735 | 0.9494 | 0.2945 | 0.5623 | **<0.0001** | **<0.0001** |
| | OB vs V2 | 0.2606 | 0.9897 | 0.9889 | 0.7840 | >0.9999 | **<0.0001** | **<0.0001** |
| | M1 vs S1 | 0.9995 | 0.6663 | 0.9999 | 0.9981 | 0.9983 | **0.0375** | 0.9171 |
| | M1 vs V2 | 0.7673 | 0.7844 | 0.6850 | **0.0442** | 0.1482 | **<0.0001** | 0.1486 |
| | S1 vs V2 | 0.9533 | 0.0676 | 0.6048 | **0.0146** | 0.4103 | **0.0016** | 0.7756 |
| **NREM** | OB vs M1 | **0.0090** | **0.0062** | **0.0124** | **0.0389** | 0.9557 | 0.9998 | 0.8640 |
| | OB vs S1 | **<0.0001** | **0.0010** | **0.0095** | **0.0030** | 0.5119 | >0.9999 | 0.9923 |
| | OB vs V2 | 0.9203 | 0.8132 | 0.4517 | 0.3883 | 0.9637 | 0.8570 | 0.9568 |
| | M1 vs S1 | **0.0468** | 0.9888 | >0.9999 | 0.9344 | 0.9664 | 0.9954 | 0.9981 |
| | M1 vs V2 | **0.0003** | **<0.0001** | **<0.0001** | **<0.0001** | >0.9999 | 0.9667 | >0.9999 |
| | S1 vs V2 | **<0.0001** | **<0.0001** | **<0.0001** | **<0.0001** | 0.9591 | 0.7301 | >0.9999 |
| **REM** | OB vs M1 | >0.9999 | 0.9977 | 0.7833 | 0.2688 | 0.1028 | 0.3289 | >0.9999 |
| | OB vs S1 | >0.9999 | **0.0027** | 0.6559 | 0.0636 | **0.0472** | 0.3550 | 0.9946 |
| | OB vs V2 | 0.9997 | 0.7065 | >0.9999 | 0.9922 | >0.9999 | 0.9576 | 0.8884 |
| | M1 vs S1 | >0.9999 | **0.0120** | >0.9999 | 0.9844 | 0.9994 | >0.9999 | 0.9626 |
| | M1 vs V2 | 0.9978 | 0.9490 | 0.5988 | 0.0685 | 0.1408 | **0.0486** | 0.7387 |
| | S1 vs V2 | 0.9993 | 0.1208 | 0.4684 | **0.0122** | 0.0664 | 0.0581 | 0.9984 |

**Supplementary Figure 1. Absolute power in function of behavioral states and cortical regions.** p values of the Sidak multiple comparisons test, comparing the differences in the absolute power between behavioral states (A) and cortical regions (B). Data show in A are summarized in Figure 3. The data are from the right hemisphere during the light phase. OB, olfactory bulb; M1, primary motor cortex; S1, primary somato-sensory cortex; V2, secondary visual cortex; W, wakefulness; LG, low gamma; HG, high gamma; HFO, high frequency oscillations.

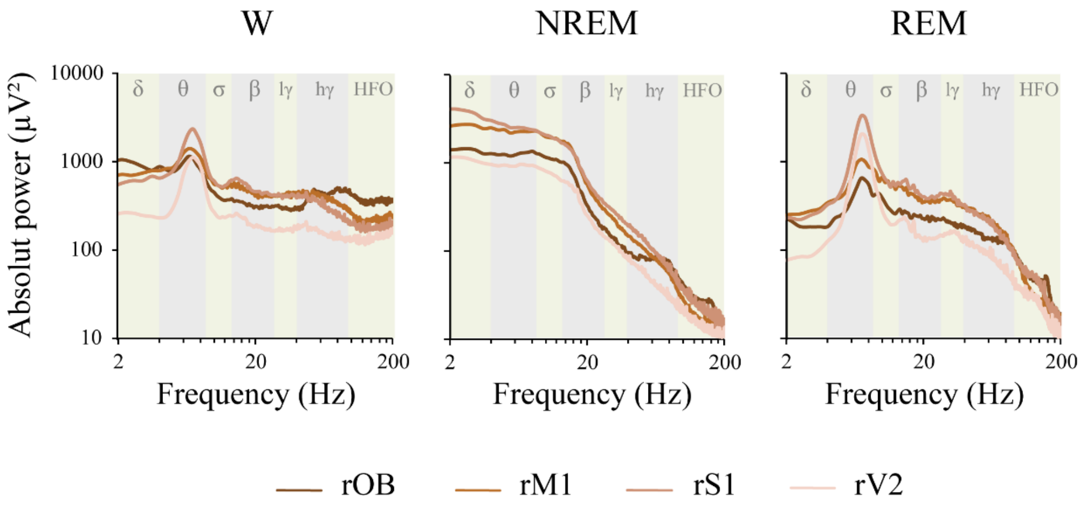


**Supplementary Figure 2. Mean absolute power in function of cortical regions.** Mean absolute power spectral profile of each behavioral state during the light period for all the electrodes of the right hemisphere. The analyzed frequency bands are indicated by different colors in the background of the graphics. OB, olfactory bulb; M1, primary motor cortex; S1, primary somato-sensory cortex; V2, secondary visual cortex; W, wakefulness; lγ, low gamma or LG; hγ, high gamma or HG; HFO, high frequency oscillations.

### Total Power

A

| Cortex | | | Behavioral State | | | Cortex x State | | |
|---|---|---|---|---|---|---|---|---|
| df | p | F | df | p | F | df | P | F |
| 3,30 | **0.0001** | 9.6 | 2, 20 | **<0.0001** | 20.0 | 6, 57 | **0.0003** | 5.2 |

B

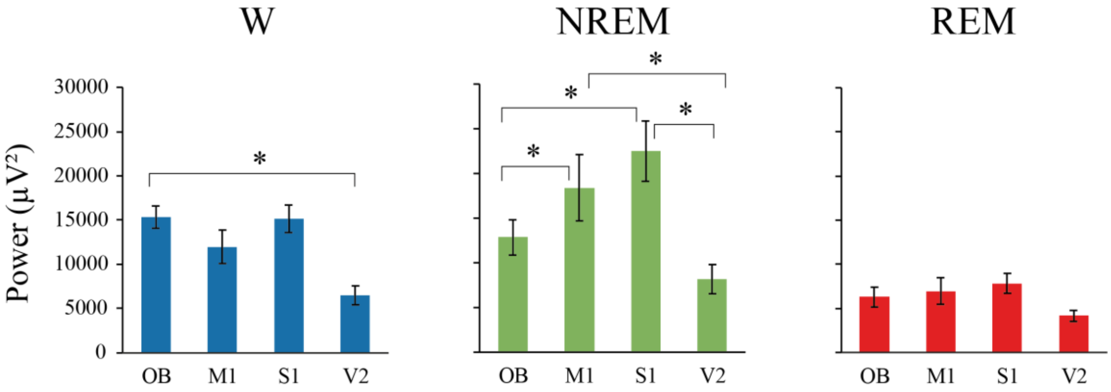


C

| State | Comparison | p value |
|---|---|---|
| **W** | OB vs M1 | 0.4853 |
| | OB vs S1 | 0.2315 |
| | OB vs V2 | 0.0005 |
| | M1 vs S1 | 0.9991 |
| | M1 vs V2 | 0.1122 |
| | S1 vs V2 | 0.2784 |
| **NREM** | OB vs M1 | **0.0163** |
| | OB vs S1 | **0.0004** |
| | OB vs V2 | 0.5393 |
| | M1 vs S1 | 0.8690 |
| | M1 vs V2 | **<0.0001** |
| | S1 vs V2 | **<0.0001** |
| **REM** | OB vs M1 | 0.8917 |
| | OB vs S1 | 0.9284 |
| | OB vs V2 | 0.9800 |
| | M1 vs S1 | >0.9999 |
| | M1 vs V2 | 0.4337 |
| | S1 vs V2 | 0.5000 |

D

| Cortex | Comparison | p value |
|---|---|---|
| **OB** | W vs NREM | 0.1968 |
| | W vs REM | **<0.0001** |
| | NREM vs REM | **0.0312** |
| **M1** | W vs NREM | **0.0144** |
| | W vs REM | 0.2855 |
| | NREM vs REM | **<0.0001** |
| **S1** | W vs NREM | **0.0001** |
| | W vs REM | 0.4712 |
| | NREM vs REM | **<0.0001** |
| **V2** | W vs NREM | 0.8431 |
| | W vs REM | 0.6690 |
| | NREM vs REM | 0.2245 |

**Supplementary Figure 3. Total power.** A. Statistical evaluation of the total power in function of cortical regions, behavioral state, and interaction between both factors. Repeated mixed-effects model. B. Mean total power in function of cortical regions. Total power is lower in the V2 probably because it is closer to the reference electrode located in the cerebellum. The error bars show the standard error of the mean. Asterisks indicate significant differences, $p < 0.05$. C. p values of the Sidak multiple comparisons test, comparing the differences in the total power between the different cortical localizations of the right hemisphere for each behavioral state. D. p values of the Sidak multiple comparisons test, comparing the differences in the total power between the different behavioral states in each right cortex. OB, olfactory bulb; M1, primary motor cortex; S1, primary somato-sensory cortex; V2, secondary visual cortex; W, wakefulness.

| Cortex | Comparison | Delta | Theta | Sigma | Beta | LG | HG | HFO |
|---|---|---|---|---|---|---|---|---|
| **OB** | W vs NREM | **0.0032** | 0.6824 | **<0.0001** | **0.0015** | **0.0314** | **<0.0001** | **<0.0001** |
| | W vs REM | 0.0926 | 0.9053 | 0.1590 | 0.8238 | **0.0068** | 0.1815 | **<0.0001** |
| | NREM vs REM | **<0.0001** | 0.9710 | **0.0134** | **0.0165** | **<0.0001** | **<0.0001** | 0.0964 |
| **M1** | W vs NREM | **<0.0001** | 0.8538 | **<0.0001** | **0.0001** | **<0.0001** | **<0.0001** | **<0.0001** |
| | W vs REM | 0.2034 | 0.8360 | **0.0066** | 0.2606 | 0.0172 | 0.7641 | **0.0002** |
| | NREM vs REM | **<0.0001** | >0.9999 | 0.3633 | **0.0272** | **<0.0001** | **<0.0001** | 0.6090 |
| **S1** | W vs NREM | **<0.0001** | **<0.0001** | **0.0195** | 0.1988 | **<0.0001** | **<0.0001** | **<0.0001** |
| | W vs REM | 0.0677 | **0.0022** | 0.1235 | 0.7243 | 0.8881 | 0.4177 | **0.0049** |
| | NREM vs REM | **<0.0001** | **<0.0001** | 0.8450 | 0.7589 | **<0.0001** | **0.0071** | 0.5565 |
| **V2** | W vs NREM | **<0.0001** | 0.7643 | **0.0003** | **0.0294** | **0.0028** | **0.0002** | **<0.0001** |
| | W vs REM | 0.1089 | **<0.0001** | 0.5729 | 0.7624 | 0.9970 | 0.1371 | **<0.0001** |
| | NREM vs REM | **<0.0001** | **<0.0001** | **0.0115** | 0.2260 | **0.0049** | **0.0861** | 0.6982 |

**Supplementary Figure 4. Differences in relative power in function of behavioral states.** p values of the Sidak multiple comparisons test, comparing the differences in the relative power in function of behavioral states for each cortical region (of the right hemisphere). Data showed in B are summarized in Figure 3. OB, olfactory bulb; M1, primary motor cortex; S1, primary somato-sensory cortex; V2, secondary visual cortex; df, degrees of freedom; W, wakefulness; LG, low gamma; HG, high gamma; HFO, high frequency oscillations.

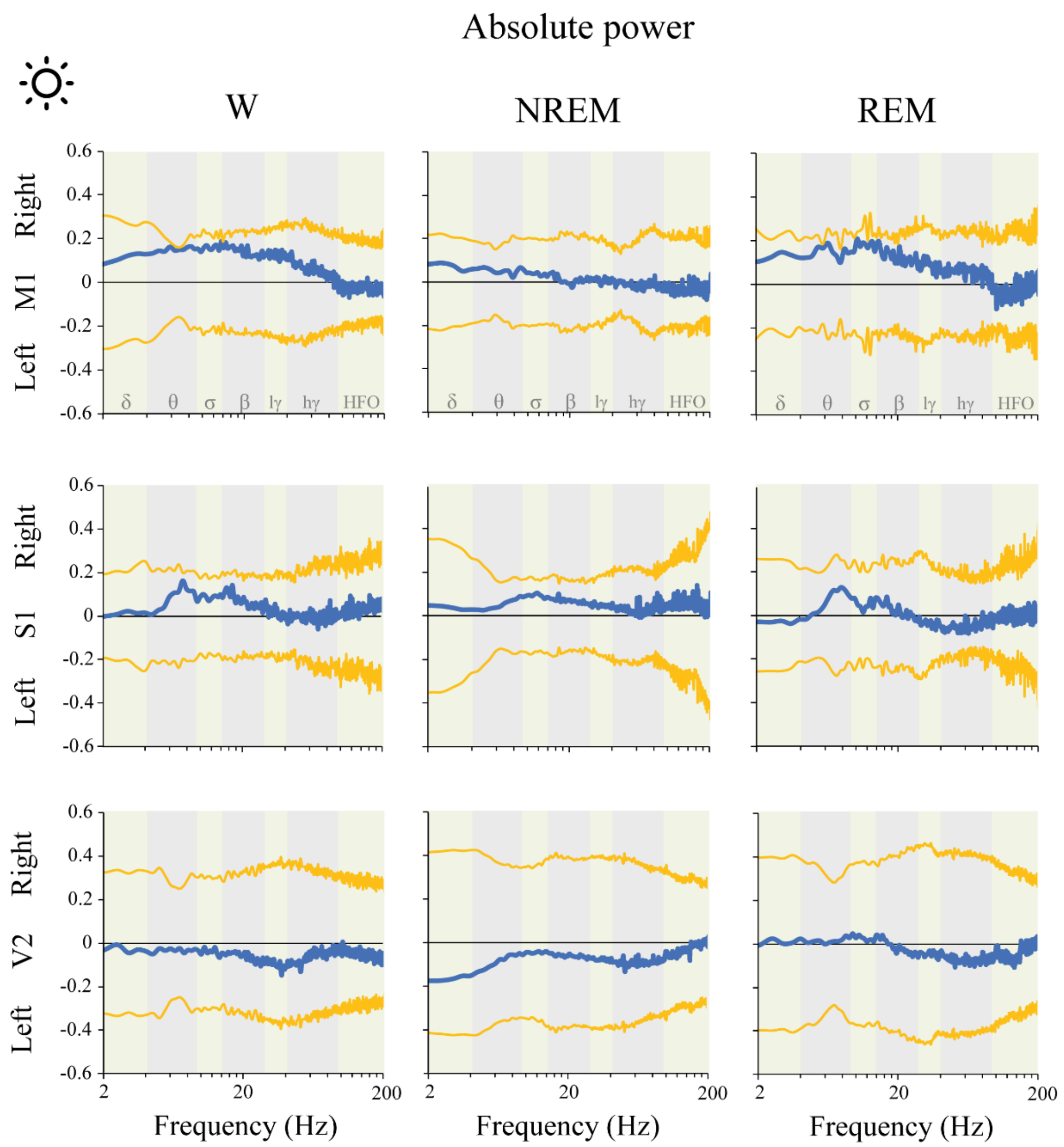


**Supplementary Figure 5. Absolute power: right Vs. left hemispheric difference during the light phase.** The predominance was calculated by means of the formula: (a-b)/(a+b). "a" represents the mean power for each frequency in the right hemisphere, and "b" the mean power in the left hemisphere. A positive value means that power in the right was higher than in the left hemisphere and *vice versa*. The blue traces indicate the mean power difference between right and dark hemispheres. The yellow lines represent the standard deviation of the mean with respect to zero. The statistical evaluation was performed by the two-tailed paired t-test with Bonferroni correction for multiple comparisons; no significant differences were observed. M1, primary motor cortex; S1, primary somato-sensory cortex; V2, secondary visual cortex; lγ, low gamma or LG; hγ, high gamma or HG; HFO, high frequency oscillations.

Absolute power

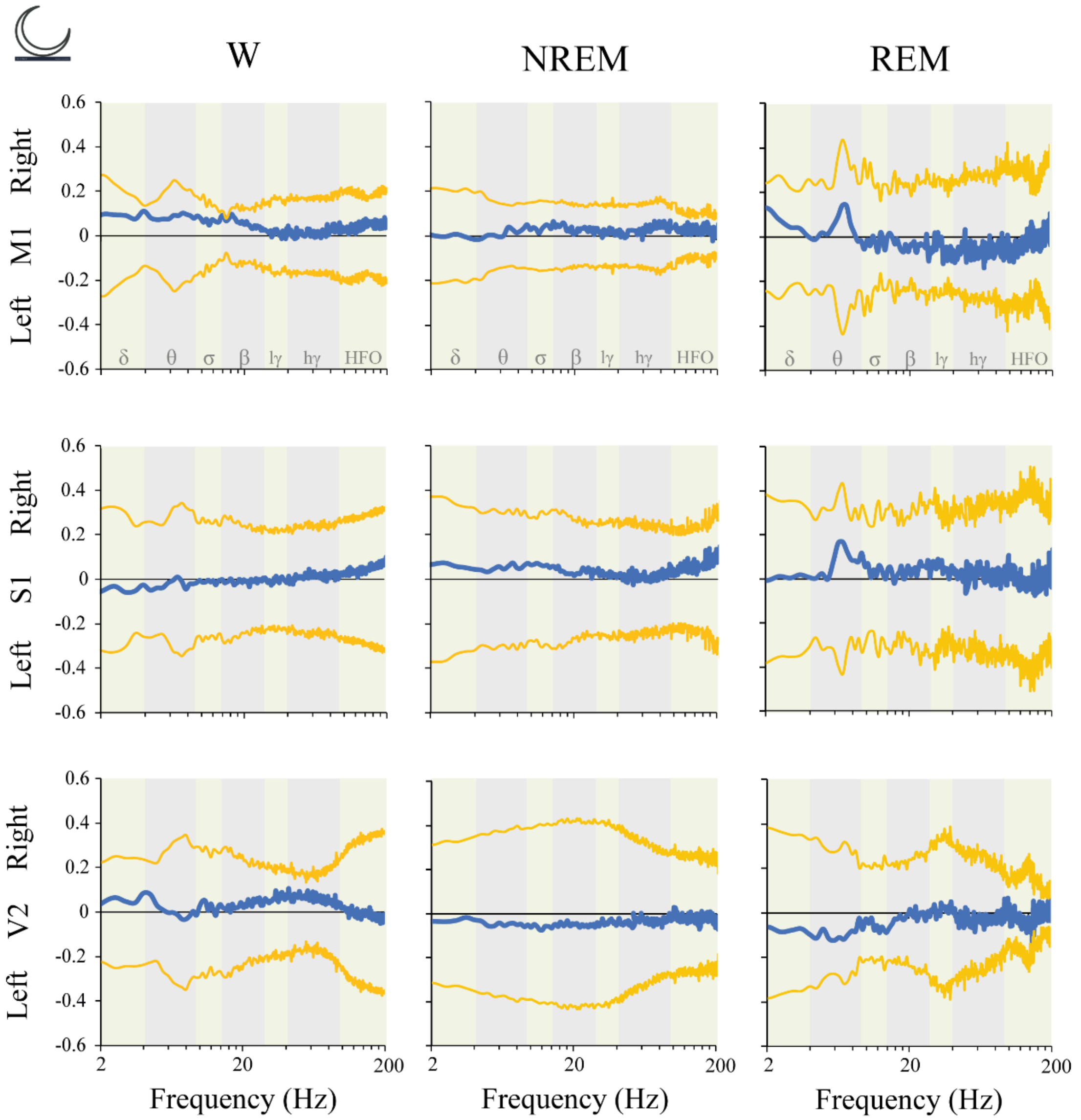


**Supplementary Figure 6. Absolute power: right Vs. left hemispheric difference during the dark phase.** The predominance was calculated by means of the formula: (a-b)/(a+b). "a" represents the mean power for each frequency in the right hemisphere, and "b" the mean power in the left hemisphere. A positive value means that power in the right was higher than in the left hemisphere and *vice versa*. The blue traces indicate the mean power difference between right and dark hemispheres. The yellow lines represent the standard deviation of the mean with respect to zero. The statistical evaluation was performed by the two-tailed paired t-test with Bonferroni correction for multiple comparisons; no significant differences were observed. M1, primary motor cortex; S1, primary somato-sensory cortex; V2, secondary visual cortex; lγ, low gamma or LG; hγ, high gamma or HG; HFO, high frequency oscillations.

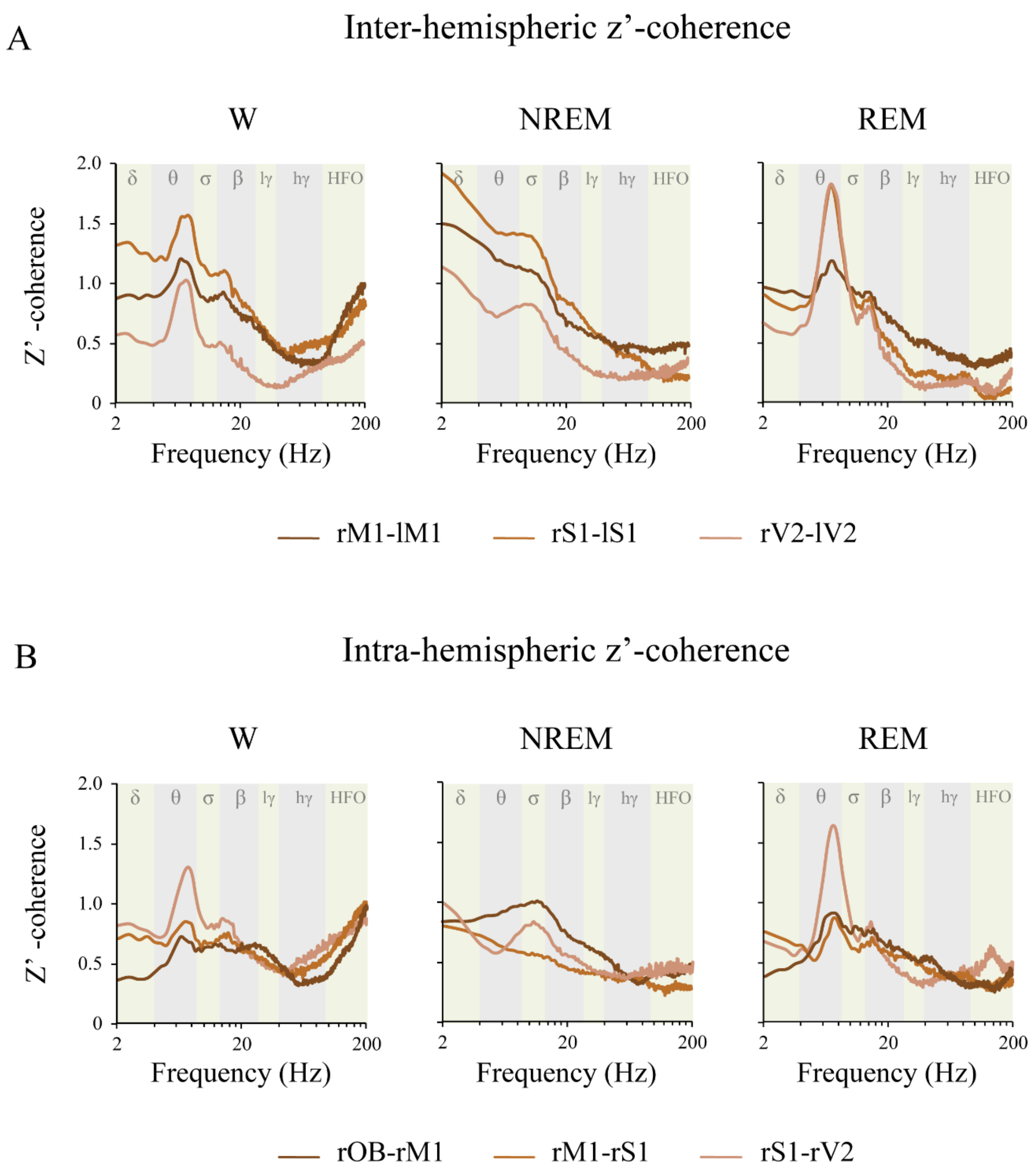


**Supplementary Figure 7. Z'-coherence in function of the derivations.** Mean z'-coherence profile of the intra-hemispheric (A, for the right hemisphere) and inter-hemispheric (B) derivations during wakefulness (W), NREM and REM sleep in the light phase. The analyzed frequency bands are indicated by different colors in the background of the graphics. OB, olfactory bulb; M1, primary motor cortex; S1, primary somato-sensory cortex; V2, secondary visual cortex; r, right; l, left; lγ, low gamma or LG; hγ, high gamma or HG; HFO, high frequency oscillations.

| Derivation | | Delta | Theta | Sigma | Beta | LG | HG | HFO |
|---|---|---|---|---|---|---|---|---|
| | Intra-hemispheric Z'-coherence | | | | | | | |
| rOB - rM1 | W vs NREM | **0.0014** | 0.0287 | **0.0004** | 0.6053 | 0.8309 | 0.9930 | **0.0007** |
| | W vs REM | 0.8569 | 0.0778 | 0.1541 | 0.9999 | 0.8453 | >0.9999 | **<0.0001** |
| | NREM vs REM | **0.0129** | 0.9520 | 0.1089 | 0.6477 | >0.9999 | 0.9935 | 0.7875 |
| rM1 - rS1 | W vs NREM | **0.0030** | 0.9974 | 0.9635 | 0.0966 | 0.8563 | 0.0898 | **<0.0001** |
| | W vs REM | >0.9999 | 0.9664 | 0.7950 | 0.6974 | 0.9471 | **0.0103** | **<0.0001** |
| | NREM vs REM | **0.0030** | 0.9192 | 0.9772 | 0.9410 | 0.9929 | 0.8696 | 0.9122 |
| rS1 - rV2 | W vs NREM | 0.9485 | **<0.0001** | 0.7063 | 0.4468 | 0.9998 | **0.0021** | **<0.0001** |
| | W vs REM | 0.4460 | 0.0731 | 0.9510 | 0.1566 | 0.1528 | **0.0004** | **<0.0001** |
| | NREM vs REM | 0.2151 | **<0.0001** | 0.9416 | 0.9410 | 0.2083 | 0.9872 | 0.9676 |
| | Inter-hemispheric Z'-coherence | | | | | | | |
| rM1 - lM1 | W vs NREM | **<0.0001** | 0.4673 | 0.0438 | 0.9834 | 0.3244 | 0.3029 | **0.0002** |
| | W vs REM | 0.9196 | 0.9992 | 0.9847 | 0.6268 | 0.9990 | 0.8330 | **<0.0001** |
| | NREM vs REM | **<0.0001** | 0.3982 | 0.0937 | 0.8552 | 0.2631 | 0.0648 | 0.3631 |
| rS1 - lS1 | W vs NREM | **0.0004** | 0.6795 | 0.8996 | 0.0922 | 0.4033 | **<0.0001** | **<0.0001** |
| | W vs REM | 0.5178 | 0.0446 | 0.7298 | **0.0011** | **0.0034** | **<0.0001** | **<0.0001** |
| | NREM vs REM | **<0.0001** | 0.0031 | 0.3474 | 0.4359 | 0.2082 | >0.9999 | 0.6549 |
| rV2 - lV2 | W vs NREM | **0.0002** | 0.6415 | **0.0062** | **0.0283** | **0.0011** | >0.9999 | 0.4611 |
| | W vs REM | 0.9918 | **<0.0001** | **0.0019** | 0.7857 | 0.9989 | **0.0322** | **0.0006** |
| | NREM vs REM | **0.0005** | **<0.0001** | 0.9944 | 0.1954 | **0.0007** | **0.0479** | 0.0567 |

**Supplementary Figure 8. Differences in z'-coherence in function of behavioral states.** p values of the Sidak multiple comparisons test, comparing the differences in the z'-coherence in function of behavioral states for each frequency band. OB, olfactory bulb; M1, primary motor cortex; S1, primary somato-sensory cortex; V2, secondary visual cortex; r, right; l, left; W, wakefulness; LG, low gamma; HG, high gamma; HFO, high frequency oscillations.

A. Wakefulness

| Comparison | Delta | Theta | Sigma | Beta | LG | HG | HFO |
|---|---|---|---|---|---|---|---|
| rOB-rM1 vs rM1-rS1 | 0.9955 | >0.9999 | >0.9999 | >0.9999 | >0.9999 | >0.9999 | 0.9998 |
| rOB-rM1 vs rM1-rV2 | 0.1567 | 0.0751 | 0.5571 | 0.9847 | 0.9947 | 0.2005 | 0.4356 |
| rOB-rM1 vs rM1-lM1 | 0.8031 | 0.8266 | 0.9969 | >0.9999 | >0.9999 | >0.9999 | >0.9999 |
| rOB-rM1 vs rS1-lS1 | **0.0235** | **0.0461** | 0.3920 | 0.9867 | >0.9999 | 0.9520 | 0.9995 |
| rOB-rM1 vs rV2-lV2 | >0.9999 | 0.9985 | >0.9999 | 0.9667 | 0.8911 | >0.9999 | 0.7259 |
| rM1-rS1 vs rS1-rV2 | 0.8703 | 0.2318 | 0.7258 | 0.9695 | 0.9755 | 0.6631 | 0.9533 |
| rM1-rS1 vs rM1-lM1 | >0.9999 | 0.9846 | 0.9998 | >0.9999 | >0.9999 | >0.9999 | >0.9999 |
| rM1-rS1 vs rS1-lS1 | 0.3692 | 0.1547 | 0.5566 | 0.9730 | >0.9999 | >0.9999 | >0.9999 |
| rM1-rS1 vs rV2-lV2 | >0.9999 | >0.9999 | >0.9999 | 0.9830 | 0.9592 | 0.9946 | 0.1782 |
| rS1-rV2 vs rM1-lM1 | 0.9986 | 0.9755 | 0.9981 | 0.9999 | 0.9926 | 0.2924 | 0.7587 |
| rS1-rV2 vs rS1-lS1 | >0.9999 | >0.9999 | >0.9999 | >0.9999 | >0.9999 | 0.9890 | 0.9727 |
| rS1-rV2 vs rV2-lV2 | 0.5609 | 0.5958 | 0.2650 | 0.2091 | 0.1661 | 0.0673 | 0.0027 |
| rM1-lM1 vs rS1-lS1 | 0.8428 | 0.9313 | 0.9860 | 0.9999 | >0.9999 | 0.9847 | >0.9999 |
| rM1-lM1 vs rV2-lV2 | 0.9960 | >0.9999 | 0.9340 | 0.7698 | 0.9079 | >0.9999 | 0.4025 |
| rS1-lS1 vs rV2-lV2 | 0.1430 | 0.4580 | 0.9963 | 0.2179 | 0.5124 | 0.7255 | 0.1447 |

B. NREM sleep

| Comparison | Delta | Theta | Sigma | Beta | LG | HG | HFO |
|---|---|---|---|---|---|---|---|
| rOB-rM1 vs rM1-rS1 | 0.9998 | >0.9999 | >0.9999 | >0.9999 | >0.9999 | >0.9999 | >0.9999 |
| rOB-rM1 vs rM1-rV2 | 0.6713 | 0.0777 | 0.9631 | 0.9998 | >0.9999 | 0.8373 | 0.3460 |
| rOB-rM1 vs rM1-lM1 | 0.8441 | 0.9997 | >0.9999 | >0.9999 | >0.9999 | >0.9999 | >0.9999 |
| rOB-rM1 vs rS1-lS1 | 0.1835 | **0.0395** | 0.9393 | >0.9999 | >0.9999 | >0.9999 | 0.9976 |
| rOB-rM1 vs rV2-lV2 | >0.9999 | 0.4186 | >0.9999 | 0.9891 | 0.8162 | 0.9974 | 0.9729 |
| rM1-rS1 vs rS1-rV2 | 0.9951 | **0.0203** | 0.6591 | 0.9928 | 0.9966 | 0.8247 | 0.1439 |
| rM1-rS1 vs rM1-lM1 | 0.9997 | 0.9701 | 0.9921 | >0.9999 | >0.9999 | >0.9999 | >0.9999 |
| rM1-rS1 vs rS1-lS1 | 0.7377 | **0.0094** | 0.5897 | >0.9999 | >0.9999 | >0.9999 | >0.9999 |
| rM1-rS1 vs rV2-lV2 | >0.9999 | 0.1581 | >0.9999 | 0.9995 | 0.9721 | 0.9979 | 0.9993 |
| rS1-rV2 vs rM1-lM1 | >0.9999 | 0.5115 | >0.9999 | >0.9999 | 0.9997 | 0.8352 | 0.2841 |
| rS1-rV2 vs rS1-lS1 | >0.9999 | >0.9999 | >0.9999 | >0.9999 | 0.9999 | 0.9349 | **0.0257** |
| rS1-rV2 vs rV2-lV2 | 0.9442 | >0.9999 | 0.9692 | 0.5786 | 0.3192 | 0.1553 | **0.0104** |
| rM1-lM1 vs rS1-lS1 | 0.9986 | 0.3355 | 0.9997 | >0.9999 | >0.9999 | >0.9999 | 0.9992 |
| rM1-lM1 vs rV2-lV2 | 0.9895 | 0.9614 | >0.9999 | 0.9549 | 0.9088 | 0.9975 | 0.9867 |
| rS1-lS1 vs rV2-lV2 | 0.4753 | 0.9985 | 0.9481 | 0.8407 | 0.8832 | 0.9835 | >0.9999 |

#### C. REM sleep

| Comparison | Delta | Theta | Sigma | Beta | LG | HG | HFO |
|---|---|---|---|---|---|---|---|
| rOB-rM1 vs rM1-rS1 | 0.9985 | >0.9999 | 0.9963 | >0.9999 | >0.9999 | >0.9999 | >0.9999 |
| rOB-rM1 vs rM1-rV2 | 0.9362 | >0.9999 | >0.9999 | >0.9999 | 0.9988 | 0.8319 | 0.8575 |
| rOB-rM1 vs rM1-lM1 | 0.2233 | 0.9882 | >0.9999 | >0.9999 | >0.9999 | >0.9999 | >0.9999 |
| rOB-rM1 vs rS1-lS1 | **0.0182** | 0.7371 | 0.9798 | >0.9999 | >0.9999 | >0.9999 | 0.9994 |
| rOB-rM1 vs rV2-lV2 | 0.9997 | >0.9999 | >0.9999 | 0.9975 | 0.9960 | >0.99990 | >0.9999 |
| rM1-rS1 vs rS1-rV2 | >0.9999 | 0.9996 | 0.8436 | 0.9275 | 0.9538 | 0.8801 | 0.4688 |
| rM1-rS1 vs rM1-lM1 | 0.8948 | 0.8642 | 0.8392 | 0.9995 | >0.9999 | >0.9999 | 0.9999 |
| rM1-rS1 vs rS1-lS1 | 0.2544 | 0.4073 | 0.3459 | 0.9755 | >0.9999 | >0.9999 | >0.9999 |
| rM1-rS1 vs rV2-lV2 | >0.9999 | >0.9999 | >0.9999 | >0.9999 | >0.9999 | >0.9999 | >0.9999 |
| rS1-rV2 vs rM1-lM1 | 0.9954 | >0.9999 | >0.9999 | >0.9999 | 0.9996 | 0.9895 | 0.9579 |
| rS1-rV2 vs rS1-lS1 | 0.5884 | 0.9619 | >0.9999 | >0.9999 | 0.9999 | 0.9125 | 0.2274 |
| rS1-rV2 vs rV2-lV2 | >0.9999 | 0.9999 | 0.9834 | 0.7859 | 0.5706 | 0.4580 | 0.5166 |
| rM1-lM1 vs rS1-lS1 | 0.9992 | >0.9999 | >0.9999 | >0.9999 | >0.9999 | >0.9999 | 0.9917 |
| rM1-lM1 vs rV2-lV2 | 0.8305 | 0.9025 | 0.9824 | 0.9910 | 0.9906 | 0.9982 | >0.9999 |
| rS1-lS1 vs rV2-lV2 | 0.1946 | 0.4658 | 0.6645 | 0.8924 | 0.9828 | >0.9999 | >0.9999 |

**Supplementary Figure 9. Z'-coherence in function of the derivation.** p values of the Sidak multiple comparisons test, comparing the differences in the z'-coherence according to the derivation, during wakefulness (A), NREM (B) and REM sleep (C). OB, olfactory bulb; M1, primary motor cortex; S1, primary somato-sensory cortex; V2, secondary visual cortex; LG, low gamma; HG, high gamma; HFO, high frequency oscillations.

| Derivation | | Delta | Theta | Sigma | Beta | LG | HG | HFO |
|---|---|---|---|---|---|---|---|---|
| | Intra-hemispheric Z'-coherence | | | | | | | |
| **rOB - rM1** | W vs NREM | **0.0014** | 0.0287 | **0.0004** | 0.6053 | 0.8309 | 0.9930 | **0.0007** |
| | W vs REM | 0.8569 | 0.0778 | 0.1541 | 0.9999 | 0.8453 | >0.9999 | **<0.0001** |
| | NREM vs REM | **0.0129** | 0.9520 | 0.1089 | 0.6477 | >0.9999 | 0.9935 | 0.7875 |
| **rM1 - rS1** | W vs NREM | **0.0030** | 0.9974 | 0.9635 | 0.0966 | 0.8563 | 0.0898 | **<0.0001** |
| | W vs REM | >0.9999 | 0.9664 | 0.7950 | 0.6974 | 0.9471 | **0.0103** | **<0.0001** |
| | NREM vs REM | **0.0030** | 0.9192 | 0.9772 | 0.9410 | 0.9929 | 0.8696 | 0.9122 |
| **rS1 - rV2** | W vs NREM | 0.9485 | **<0.0001** | 0.7063 | 0.4468 | 0.9998 | **0.0021** | **<0.0001** |
| | W vs REM | 0.4460 | 0.0731 | 0.9510 | 0.1566 | 0.1528 | **0.0004** | **<0.0001** |
| | NREM vs REM | 0.2151 | **<0.0001** | 0.9416 | 0.9410 | 0.2083 | 0.9872 | 0.9676 |
| | Inter-hemispheric Z'-coherence | | | | | | | |
| **rM1 - lM1** | W vs NREM | **<0.0001** | 0.4673 | 0.0438 | 0.9834 | 0.3244 | 0.3029 | **0.0002** |
| | W vs REM | 0.9196 | 0.9992 | 0.9847 | 0.6268 | 0.9990 | 0.8330 | **<0.0001** |
| | NREM vs REM | **<0.0001** | 0.3982 | 0.0937 | 0.8552 | 0.2631 | 0.0648 | 0.3631 |
| **rS1 - lS1** | W vs NREM | **0.0004** | 0.6795 | 0.8996 | 0.0922 | 0.4033 | **<0.0001** | **<0.0001** |
| | W vs REM | 0.5178 | 0.0446 | 0.7298 | **0.0011** | **0.0034** | **<0.0001** | **<0.0001** |
| | NREM vs REM | **<0.0001** | 0.0031 | 0.3474 | 0.4359 | 0.2082 | >0.9999 | 0.6549 |
| **rV2 - lV2** | W vs NREM | **0.0002** | 0.6415 | **0.0062** | **0.0283** | **0.0011** | >0.9999 | 0.4611 |
| | W vs REM | 0.9918 | **<0.0001** | **0.0019** | 0.7857 | 0.9989 | **0.0322** | **0.0006** |
| | NREM vs REM | **0.0005** | **<0.0001** | 0.9944 | 0.1954 | **0.0007** | **0.0479** | 0.0567 |

**Supplementary Figure 10. Differences in z'-coherence in function of behavioral states.** p values of the Sidak multiple comparisons test, comparing the differences in the z'-coherence between behavioral states for each frequency band. OB, olfactory bulb; M1, primary motor cortex; S1, primary somato-sensory cortex; V2, secondary visual cortex; r, right; l, left; W, wakefulness; LG, low gamma; HG, high gamma; HFO, high frequency oscillations.

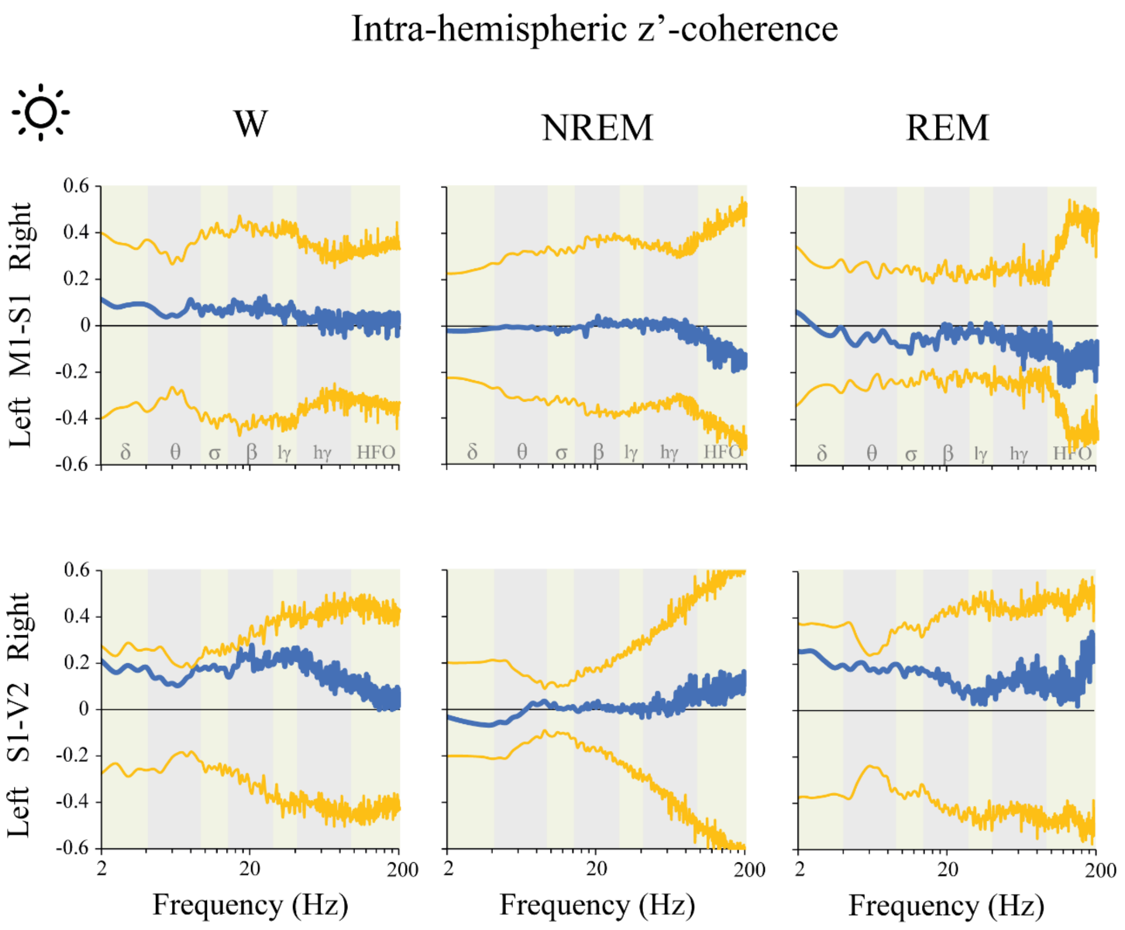


**Supplementary Figure 11. Intra-hemispheric z'-coherence: right Vs. left hemispheric difference during the light phase.** The predominance was calculated by means of the formula: (a-b)/(a+b). "a" represents the mean z'-coherence for each frequency in the light phase, and "b" the mean coherence during the dark period. A positive value means that z'-coherence in the light period was higher than during dark period and *vice versa*. The blue traces indicate the mean z'-coherence difference between light and dark phases. The yellow lines represent the standard deviation of the mean with respect to zero. The statistical evaluation was performed by the two-tailed paired t-test with Bonferroni correction for multiple comparisons; no significant differences were observed. M1, primary motor cortex; S1, primary somato-sensory cortex; V2, secondary visual cortex; lγ, low gamma or LG; hγ, high gamma or HG; HFO, high frequency oscillations.
